# Enhancing detectable fluorescence fluctuation for high-throughput and four-dimensional live-cell super-resolution imaging

**DOI:** 10.1101/2022.12.12.520072

**Authors:** Weisong Zhao, Shiqun Zhao, Zhenqian Han, Xiangyan Ding, Guangwei Hu, Xinwei Wang, Heng Mao, Yaming Jiu, Ying Hu, Jiubin Tan, Xumin Ding, Changliang Guo, Liangyi Chen, Haoyu Li

**Author notes:** These authors contributed equally to this work (Weisong Zhao, Shiqun Zhao, Zhenqian Han, Xiangyan Ding).

## Abstract

Super-resolution (SR) imaging with high-throughput is invaluable to fast and high-precision profiling in a wide range of biomedical applications. However, prevalent SR methods require sophisticated acquisition devices and specific imaging control, and may cost a fairly long time on a single field-of-view. These essentially increase the construction difficulty, including challenges in imaging throughput, system establishment, and automation. Using the natural photophysics of fluorescence, fluctuation-based microscopy techniques can routinely break the diffraction limit with no need for additional optical components, but its long acquisition time still poses a challenge for high-throughput imaging or visualizing transient organelle dynamics. Here, we propose an <u>S</u>R method based on the <u>A</u>uto-<u>C</u>orrelation with two-step <u>D</u>econvolution (SACD) that reduces the number of frames required by maximizing the detectable fluorescence fluctuation behavior in each measurement, with further removal of tunable parameters by a Fourier ring correlation analysis. It only needs 20 frames for twofold lateral and axial resolution improvements, while the SR optical fluctuation imaging (SOFI) needs more than 1000 frames. By capturing raw images for ∼10 minutes, we record an SR image with ∼128 nm resolution that contains 2.4 gigapixels covering an area of ∼2.0 mm × 1.4 mm, including more than 2,000 cells. Beyond that, by applying continuity and sparsity joint constraint, the Sparse deconvolution-assisted SACD enables 4D live-cell SR imaging of events such as mitochondrial fission and fusion. Overall, as an open-sourced module, we anticipate SACD can offer direct access to SR, which may facilitate the biology studies of cells and organisms with high-throughput and low-cost.

---

In quantitative biology, the microscopy-based high-throughput screening is used to monitor the variability of biological systems as well as examine the heterogeneity^1^, and the advances in three-dimensional (3D) resolution can contribute to the minimization of the uncertainties in analyzing noisy biological processes. In parallel, super-resolution (SR) microscopy techniques relying on the blinking of single fluorophores have been developed to break the diffraction limit, including photoactivated localization microscopy (PALM)^2^ and stochastic optical reconstruction microscopy (STORM)^3^. However, precisely localizing individual fluorophores requires tens of thousands of frames to accumulate one final SR view, which inherently limits the throughput and leads to requirements for purpose-built experimental settings^4^. Therefore, although live-cell PALM/STORM has been reported^5-7^, excessive illumination power (∼10 kW/cm^2^)^8^, long exposures (>2 s)^9^, and the particular photochemical environment that promotes long dark states to reduce bleaching^9^ prevent it from being a general live-cell imaging method^10^ with high-throughput.

This localization mechanism essentially restricts the maximum density of emitting fluorophores in one frame^11^. Thus, it motivates techniques that rely on independent stochastic blinking/fluctuation of many emitters rather than single ones, namely fluorescence fluctuations-based SR microscopy techniques^12-15^. Remarkably, without the need for sophisticated acquisition devices, the SR optical fluctuation imaging (SOFI)^12^ calculates the *n*-th order auto-correlation cumulants yielding fold resolution improvements in all 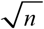 three dimensions. Because of the relaxation on using blinks of single molecules and stringent photochemical environment^16, 17^, SOFI is faster than PALM/STORM and suitable for a range of live-cell SR imaging applications, such as observing the dynamics focal adhesions^18^, clathrin-coated pits (CCPs)^19^, and actin^20^. Nevertheless, considering at least consecutive hundreds to thousands of frames are needed to reconstruct one SR image^18^, its temporal resolution and throughput are still limited.

To achieve frame-minimalism for fluctuation-based modalities, we designed an <u>S</u>R method based on <u>A</u>uto-<u>C</u>orrelation with two-step <u>D</u>econvolution (SACD). By maximizing the detectable fluorescence fluctuation behavior in each measurement, our SACD only needs 20 frames to form a high-fidelity super-resolution image achieving twofold lateral and axial resolution improvements, and the recording quality is consistent with the corresponding SOFI image which is produced by hundreds of frames. Furthermore, to achieve fully automatic calculation, we used the Fourier ring correlation (FRC)^21, 22^ to determine the iteration times of deconvolution non-biasedly. We applied SACD to directly achieve high-throughput SR imaging, in which it enables a stable SR of ∼128 nm, over a ∼2 mm × 1.4 mm area within ∼10 minutes, providing a throughput hard to achieve previously^23^. Beyond that, during long-term live-cell SR imaging, photobleaching and phototoxicity cause deterioration of signal-to-noise ratio (SNR) and cell status, which confers the fundamental limit^24^. We also adapted SACD to image reconstruction under low SNR conditions using the previously developed deconvolution solution constrained by sparsity and continuity prior knowledge^25^ in the post-deconvolution step. We could capture rapid dynamics of mitochondrial membrane structures, such as fission and fusion, up to four dimensions in the live cells.

Taken together, we anticipate that our SACD may extend the boundaries of 4D live-cell applications, allowing SR imaging of processes with high temporal resolution and long recording time. To facilitate its dissemination and future applications, we have provided the corresponding easy-to-use library and Fiji/ImageJ plugin, offering an add-on SR feature on any *off-the-shelf* commercialized microscopes or *in-house* customized systems.

## RESULTS

### SACD model development

Despite SOFI has theoretically unlimited spatial resolution, three fundamental issues limit its high-order (*n* > 2) achievement in practice: (i) nonlinear response to brightness and blinking heterogeneities^20^, (ii) potential reconstruction artifacts^26^, and (iii) the requirement of long sequences^18^. Reconstruction from extended image sequences often leads to motion artifacts blurred by highly mobile subcellular structures in live cells. Here, we focused on the second-order (*n* = 2) SOFI over the higher-order one.

In principle, the auto-correlation cumulant can eliminate any noise uncorrelated over time^12^. However, the fact is that the statistical uncertainty of auto-correlation cumulant from limited datasets may dramatically affect the continuity and homogeneity^27^. Hence, SOFI generally requires at least hundreds of raw images to preserve the structural integrity and sample details. Predictably, the magnitude of the statistical uncertainty is highly related to the quantity and accuracy of the raw data. When considering the accuracy only, two properties of each measurement influence the statistical error magnitude, including (i) the captured effective on/off contrast ratio and (ii) the SNR condition^17, 19^. In the original SOFI model, the sample is seen as being composed of *N* independently fluctuating emitters (**Extended Data Fig. 1a, Methods**)^12^. Besides in-focus fluorescent molecules, the experimental image is corrupted by out-of-focus and cytosol fluorescence signals that may not stay in stationary equilibrium (**Extended Data Fig. 1a, Methods**). These aberrant fluorescence signals compromise the effective on/off contrast ratio, while the sensor’s readout noise, dark current, and shot noise reduced the image SNR. Thus, more raw frames are required for increasing the accuracy and quality of SR images^12, 18^, which constitutes the ultimate temporal limit to fluctuation-based SR microscopy.

In this work, we applied the Richardson-Lucy (RL) deconvolution^28, 29^ on the raw image stack as a preprocessing step (**Fig. 1a**), aiming to filter the out-of-focus and cytosol background before calculating the auto-correlation cumulant. Such a procedure improved the effective on/off contrast ratio and reduced the random pixel-level fluctuations induced by noise, while maintaining the linear responses across frames (**Extended Data Fig. 2**, details in **Supplementary Note 1**). To avoid destroying the independence of fluorescence fluctuation due to excessive RL deconvolution, we cautiously stopped the iteration at an early stage. This pre-processing step significantly highlighted the structures in raw images, which effectively increased the accuracy of each measurement. Based on this preprocessing, 20 frames are workable for the next auto-correlation cumulant calculation with a Fourier interpolation^30^ (**Fig. 1a, Methods**). Lastly, we added a postprocessing step once again to deconvolve the second-order SOFI images (**Fig. 1a**), for further extending resolutions.

**Fig. 1.**
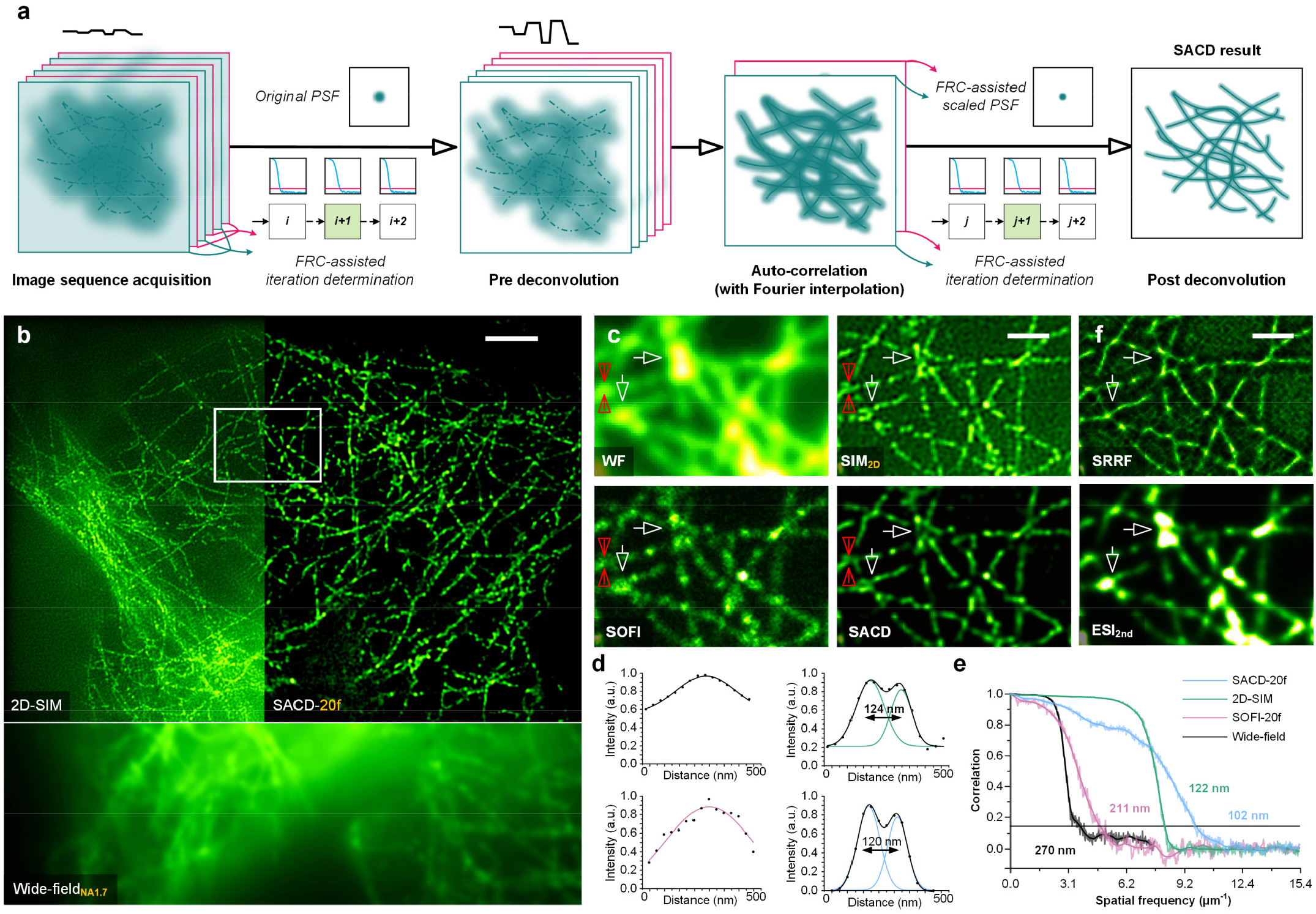
Workflow and cross-validation of SACD. (**a**) Reconstruction workflow of SACD. (**b**) Microtubule filaments in a COS-7 cell labeled with QD_525_ imaged by wide-field microscopy (bottom), 2D-SIM (left), and 20-frame SACD (right). (**c**) Magnified views from the white box in (**b**). Top panel: Wide-field image (left, averaged by 20 frames); 2D-SIM (right). Bottom panel: 20-frame SOFI (left); 20-frame SACD (right). (**d**) Intensity profiles and multiple Gaussian fitting of raw wide-field, 2D-SIM, 20-frame SOFI, and 20-frame SACD for the structures of microtubule filaments indicated by the red arrows in (**c**). The numbers indicate the distance between peaks. (**e**) FRC analysis of the reconstructed images. (**f**) Magnified views from the white box in (**b**) of 20-frame SRRF with temporal radiality average (top) and 20-frame 2nd order ESI (bottom). Scale bars: (**b**) 3 μm; (**c, f**) 1 μm.

Furthermore, to automatically adjust the tunable parameters, we introduced Fourier ring correlation (FRC)^21, 22^ as a quantitative resolution metric to assist in the process (**Extended Data Fig. 1b, Methods**). Specifically, the FRC was used to determine the optimal iteration times and estimate the effective point-spread-function (PSF) after the auto-correlation cumulant calculation^22^. Although the maximum resolution (lowest FRC value) related iteration times could be used (18 iterations in **Extended Data Fig. 3**), we intended to be conservative in choosing optimal iteration numbers for pre-deconvolution to avoid the possible disturbance on later auto-correlation. As the representative example in **Extended Data Fig. 1b**, we generally stopped the pre-RL-deconvolution at its half iteration times of maximal resolution (9 iterations in **Extended Data Fig. 3**). On the other hand, with no such concern in the post-deconvolution, the iterative process could be stopped at the maximum FRC resolution reached.

We systematically compared the performances of SACD against other state-of-the-art fluctuation-based methods in simulation (**Methods**). First, in a simple case of molecule-pair with Poisson noise only, all SOFI (20-frame and 1000-frame) and SACD (20-frame) results achieved comparable performances in high quality (**Extended Data Fig. 4b**). Then, to examine the capacity during the accuracy decreasing, we started to increase the noise levels, which were classified as ‘*high*’, ‘*medium*’, and ‘*low*’ imaging conditions (**Methods, Extended Data Fig. 4c-4d**). Upon decreases in *SNR* and *contrast*, qualities of 20-frame SOFI reconstructions continued to deteriorate while the 1000-frame one performed relatively stable. In comparison, SACD is insensitive to the *SNR* and *contrast* conditions and can effectively reconstruct high-fidelity SR images with minimum input. Next, to further demonstrate its superiority, we created more complex structures, i.e., rings with different diameters, and compared SACD against other advanced SOFI reconstruction methods^31-33^ (**Methods, Extended Data Fig. 5, Supplementary Note 2**). Despite incorporating a wavelet-based pre-filter, an RL post-deconvolution, or a different statistics calculation, the SOFI-wavelet (**Extended Data Fig. 5c**), SOFI-RL (**Extended Data Fig. 5d**), and RD-Covar (**Extended Data Fig. 5e**) exhibited dramatically decreases of image fidelity with the deterioration of imaging conditions, while the SACD (**Extended Data Fig. 5f**) consistently resolved ring structures without amplifications of the artifacts. We found that SACD could reconstruct the 180 nm ring with high-contrast, which is quantitatively determined by the ring quality ratio (RQR, **Methods, Extended Data Fig. 5h**) and equal to the ∼110 nm resolution ground truth (**Extended Data Fig. 5a**), and in contrast a ring diameter must be larger than 220 nm before distinguished by other SOFI methods. Furthermore, to evaluate the performances of different methods across different numbers of frames, we included the structural similarity (SSIM)^34^ (**Extended Data Fig. 5g**) and standard deviation (STD)^18^ (**Extended Data Fig. 5i**), reflecting the reconstruction quality and uncertainty. We found rapid increases in the fidelity and stability of SACD reconstructions before 20 frames, and after that, the SSIM and STD values reached a plateau. In contrast, SSIM and STD values of the other methods continued to vary as the number of frames rose, and did not stop even by the sequence reaching 500 frames. Likewise, both visual and quantitative examinations in the case of filament structures demonstrate the SACD can decode SR details from low contrast fluctuations with minimum input (**Supplementary Fig. 6**, details in **Supplementary Note 2**). Through these tests, we uncovered that the measured accuracy in each frame is the essential factor for fluctuation-based SR microscopy, rather than the quantity considered previously. In addition, we also replaced the pre-RL-deconvolution with different pre-processing operations, and they were not comparable to our current solution (**Supplementary Note 3**).

### Cross-validation using SIM and other fluctuation-based techniques

We first tested the performances of SOFI and SACD on a spinning disk confocal (SD-confocal) microscope (**Methods**), in which SACD consistently achieved superior image quality and resolution enhancement against the 1000-frame SOFI (**Extended Data Fig. 6**, details in **Supplementary Note 4**). Then, we intended to test the reliability of SACD in resolving fine structure details and its broad applicability to different imaging modalities. Here, a custom-built structured illumination microscope (SIM) (**Methods**)^35^ was employed to acquire 2D-SIM images of quantum dot 525 (QD_525_) labeled microtubules. The reason of choosing SIM images for cross-validation is that SIM also provides a resolution doubling of the diffraction limit (left panel of **Fig. 1b**), and in practice 2D-SIM can provide reference images *in situ* for SACD (right panel of **Fig. 1b**) which uses the diffraction-limited image sequence under equivalent wide-field illumination (**Methods**, bottom panel of **Fig. 1b**). As highlighted in **Fig. 1c**, we found the 20-frame SACD successfully resolved the intertwining filaments structures identical to the corresponding 2D-SIM image, and the 1000-frame SACD exhibited an almost identical result (**Extended Data Fig. 7c**), indicating the 20-frame input is already sufficient for our method. Benefiting from the improved axial resolution, SACD eliminated the out-of-focus light offering a higher contrast than 2D-SIM (**Fig. 1b**). Analysis of peak-to-peak separations of adjacent microtubule filaments indicated the achievable resolution of ∼120 nm for SACD, as well as ∼124 nm for 2D-SIM (**Fig. 1d**). The resolutions in the wide-field (270 nm), SOFI (211 nm), SIM (122 nm), and SACD (102 nm) images were further confirmed using the FRC analysis (**Fig. 1e**), substantially demonstrating the resolvability improvement of SACD.

Furthermore, as seen in **Fig. 1f**, taking advantage of the SIM image as a dependable reference, we also benchmarked the superior performances of SACD with three other fluctuation-based reconstruction methods, i.e., SOFI, super-resolution radial fluctuations (SRRF)^13^, and entropy-based super-resolution imaging (ESI)^15^. As pointed out by the white arrows, SOFI, SRRF, and ESI failed to resolve fine structures due to the lack of raw images. In SACD and SIM images, they could dissect such intertwining filaments, in which SACD also exhibited its superior capability for background-free and contrast-enhanced. Besides, we also compared the modified SOFI methods using the same dataset, and these methods achieved acceptable reconstruction only under the 1000-frame configuration (**Extended Data Fig. 7a-7c**, details in **Supplementary Note 7**).

### 3D resolution doubling by SACD

To adequately evaluate the resolution enhancement capability of SACD in all three dimensions, we further extended the experiments to volumetric imaging. Preliminarily, we measured its 3D PSF, in which a sample of QD_525_ was randomly dispersed on a substrate and captured by a standard SD-confocal microscope (**Fig. 2a, Methods**). We extracted the lateral/axial full-width at half-maximums (FWHMs) of this point source as 325/655 nm, 296/510 nm in SOFI, and 135/334 nm in SACD (all by 20 frames), i.e., averaged for SD-confocal and reconstructed for SOFI and SACD. The resulting FWHMs indicated that SACD effectively achieved a twofold resolution enhancement in all three dimensions (**Supplementary Fig. 1**). Compared to SOFI, the SACD effectively improved the homogeneity of PSF even using a limited dataset (20 frames).

**Fig. 2.**
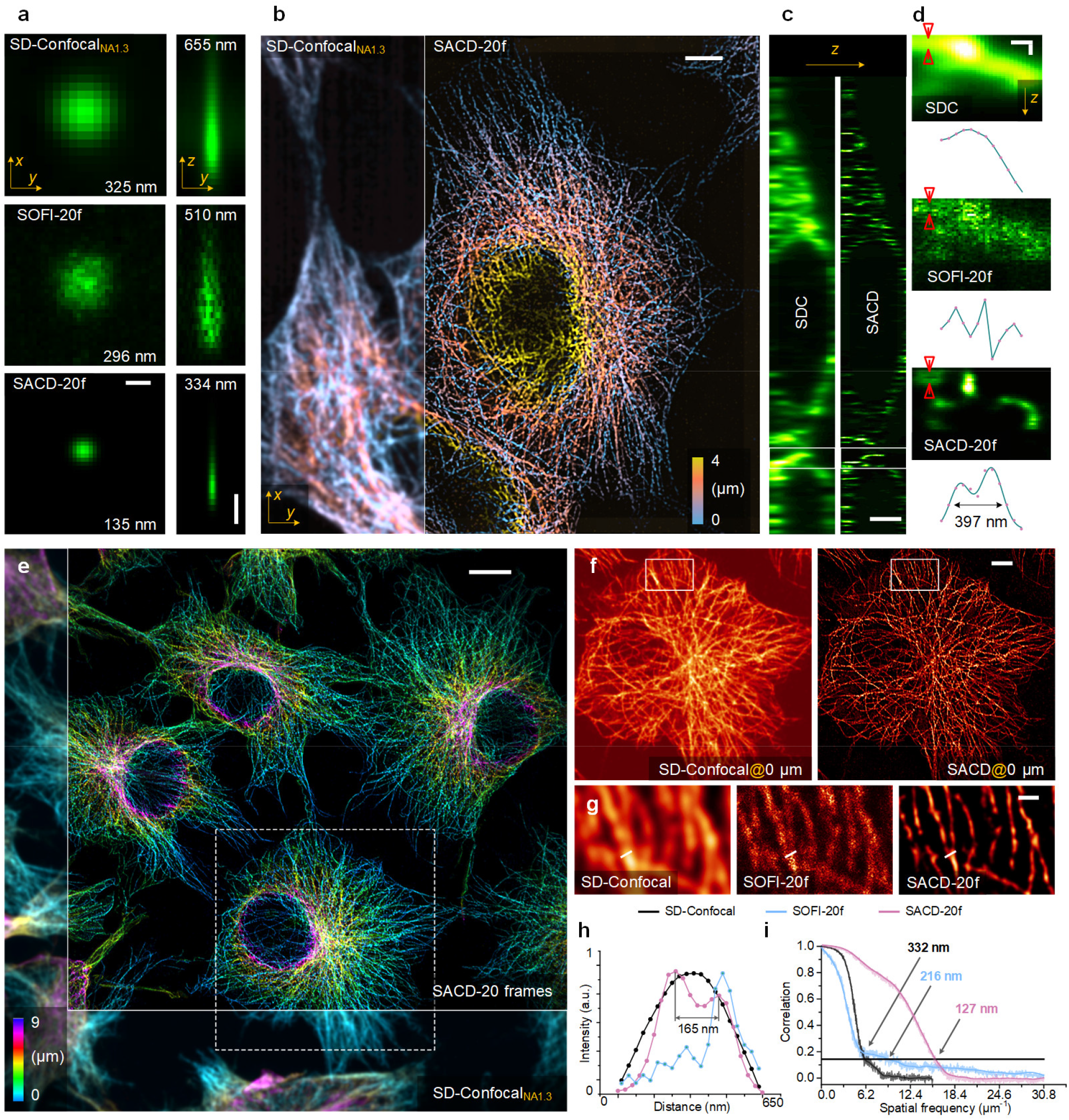
Validation for the 3D resolution doubling of SACD and large FOV volumetric imaging. (**a**) From top to bottom: The maximum intensity projections (left) and vertical sections (right) of a single QD_525_ imaged by SD-confocal (top), SOFI with 20 frames (middle), and SACD with 20 frames. (**b**) Color-coded three-dimensional distributions of microtubule filaments in a COS-7 cell labeled with QD_525_ under the SD-confocal (bottom) and SACD (top). (**c**) Corresponding vertical sections of SD-confocal (left) and SACD (right). (**d**) Axial resolution comparison using the enlarged regions of the white boxes indicated in (**c**). From left to top: cross-sections of SD-confocal, SOFI (20 frames), and SACD (20 frames) results and the intensity profiles indicated by red arrows are displayed under the corresponding configurations. (**e**) Color-coded, three-dimensional distributions of microtubule filaments in COS-7 cells labeled with QD_525_ under the SD-confocal (bottom) and SACD (top) (**Supplementary Video 1**). (**f**) Magnified horizontal section views from white dashed box in (**a**). (**g**) Enlarged regions from white dashed boxes in (**f**). (**h**) Intensity profiles of raw SD-confocal (green), 20-frame SOFI (blue), and 20-frame SACD (red) for the structures of microtubule filaments indicated by the white lines in (**g**). The numbers indicate the distance between peaks. (**i**) FRC analysis of the reconstructed images. Scale bars: (**a**, lateral and axial in **d**) 500 nm; (**b, f**) 5 μm; (**c**) 2 μm; (**e**) 10 μm; (**g**) 1 μm.

Next, we intended to exhibit the veritable 3D SR ability of SACD by imaging the microtubule cytoskeleton (**Fig. 2b, Extended Data Fig. 8**). Presenting axial cross-sections of SD-confocal (SDC) and SACD in **Fig. 2c** and **Extended Data Fig. 8**, as expected from a conventional SD-confocal microscope, we found the fine features of microtubules are blurred in all directions (left panel), especially along the axial direction. In contrast, the SACD clearly revealed the microtubule network in all three dimensions (right panel). Furthermore, focusing on the regions between red arrows in **Fig. 2d**, SACD can dissect the overlapping microtubule filaments that were completely unresolvable in conventional SD-confocal images, and the SOFI image with insufficient frames also failed to distinguish such fine structures.

Since the SACD requires no specific optical illumination and configurations, we can directly use the full sCMOS camera field-of-view (FOV >130 × 130 µm^2^, **Fig. 2e, Supplementary Fig. 2, Supplementary Video 1**) to implement 3D SR imaging. On the other hand, the existing commercial SIM setups are mostly limited to FOV with a third to a quarter of the camera (30 × 30 µm^2^ to 60 × 60 µm^2^ as shown in the white dashed box in **Fig. 2e**)^36^. Taking advantage of that, we simultaneously imaged multiple COS-7 cells at a 9 µm depth, with the doubled resolutions (**Fig. 2f-2i**). Likewise, under this configuration, we found that the 20-frame SOFI image suffered from limited quality and its continuity appeared undesirable (**Fig. 2g**). In comparison, the 20-frame SACD (**Fig. 2g**) consistently performed a reliable resolution enhancement, in which a 165 nm distance of microtubule filaments has been distinctly distinguished as stated in **Fig. 2h**. Here the FRC analysis (**Fig. 2i**) also revealed that SACD (127 nm in red) also significantly outperformed the 20-frame SOFI (216 nm in blue), and such superiority emerged in all spatial frequencies.

### Direct high-throughput SR imaging

With no need for additional optical devices or specific photoactivation control, our SACD can directly enable an add-on SR feature for various fluorescence imaging systems. Beyond that, the drastic improvement in the efficiency of SACD permits SR imaging of orders of magnitude more cells and FOVs per unit time, spontaneously benefitting high-throughput imaging. In experiments, we used an automated acquisition protocol in a commercialized SD-confocal microscope to image microtubules in a millimeter-level (2.0 mm × 1.4 mm) area with 32 × 22 partly overlapping FOVs (66.6 μm × 66.6 μm each, detailed in **Methods**). The imaging time was only within ∼10 minutes by SACD, and in contrast, the SOFI may need ∼8 hours to achieve a similar imaging performance.

As shown in **Fig. 3a** and **Supplementary Video 2**, we obtained 704 sub-FOVs in total, and each sub-FOV was captured with 20 frames. Then we stitched all resulting super-resolved sub-FOVs into one single large FOV, containing >2000 cells. Stitching the reconstructed images together yields a single SR image that contains approximately 2.4 gigapixels (32.5 nm pixel size), thereby spanning almost five orders of magnitude in spatial scales. Zooming into the randomly chosen sub-FOV, SACD images exhibit the microtubule filament structures clearly (**Fig. 3d**), and neither the SD-confocal images nor the 20-frame SOFI provided the small-scale information (**Fig. 3e, Supplementary Fig. 3a-3c**).

**Fig. 3.**
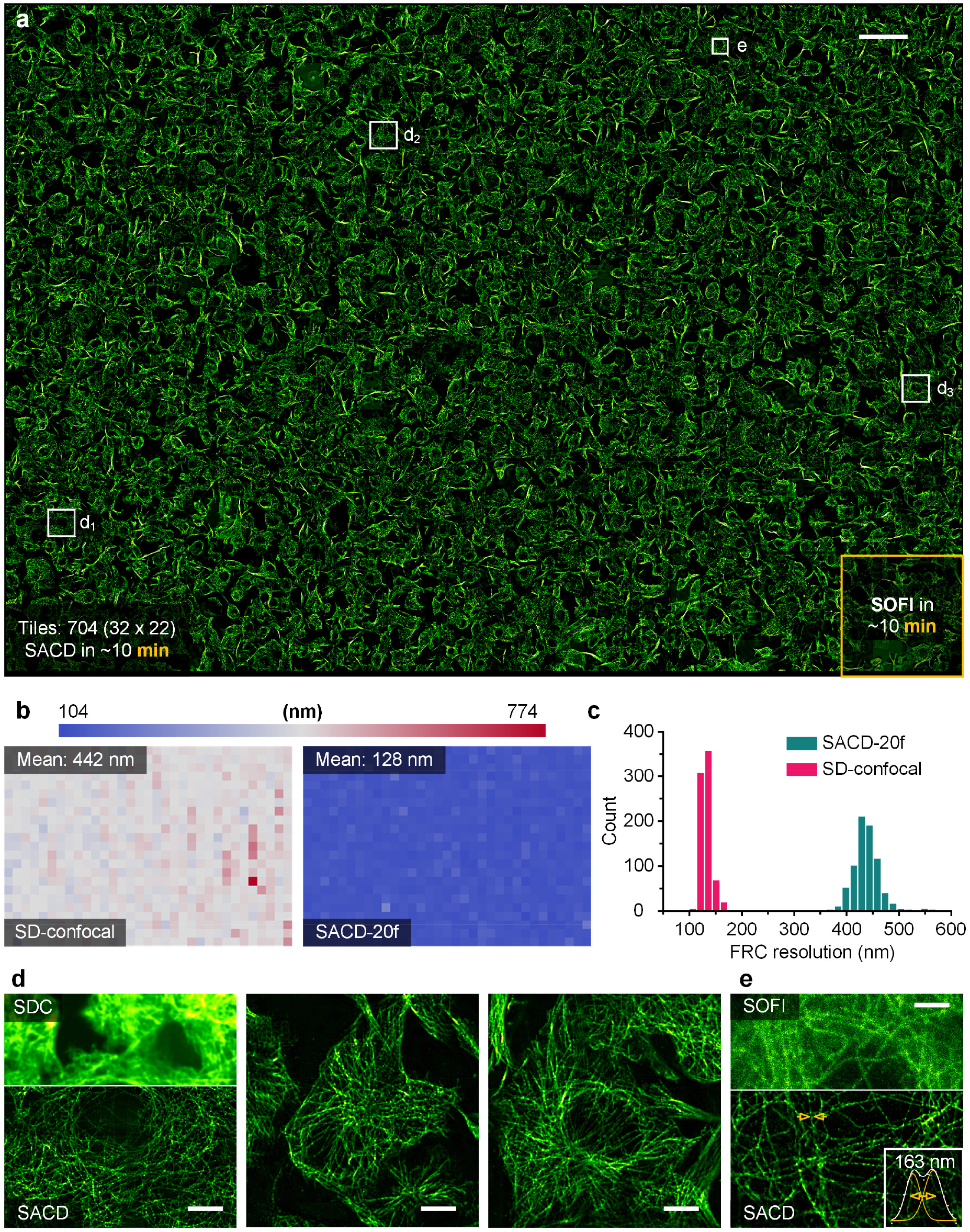
High-throughput SR imaging with SACD. (**a**) Application of SACD (with 20 frames) to high-throughput super-resolution imaging of a ∼2.0 mm × 1.4 mm area containing more than 2,000 cells. Microtubules in COS-7 cells labeled with QD_605_ (**Supplementary Video 2**). (**b**) FRC resolution maps over the entire field-of-view of SD-confocal image (left) and SACD image (right). (**c**) FRC resolution distributions of SD-confocal (magenta) and SACD (green). (**d**) Magnified views of the white boxed regions in (**a**). SDC: SD-confocal. (**e**) Enlarged view of the white boxed region in (**a**). Inset is the intensity profiles and multiple Gaussian fitting of SACD for the structures of microtubule filaments indicated by the yellow arrows in (**c**). Scale bars: (**a**) 100 μm; (**d**) 10 μm; (**e**) 5 μm.

To quantify the resolution improvements over the entire area, we measured the FRC resolutions of SD-confocal and SACD for each sub-FOV, and assembled them into a resolution map (**Fig. 3b**). SACD consistently improved the effective resolution of the SD-confocal from ∼442 nm to ∼128 nm (**Fig. 3c, Supplementary Fig. 3d**) over the entire large FOV in **Fig. 3a**. In addition, the conventional two-peak analysis also met the FRC measurement, in which a 163 nm distance of microtubule filaments were distinguished distinctly (inset of **Fig. 3e**). Ultimately, all these visualizations and analyses endorse the ability of our SACD in terms of enabling toilless high-throughput SR imaging.

In addition, with optimized detectable fluctuation behavior, the SACD concept also shows its flexibility, in which it can be used to combine with other widely adopted fluctuation-based techniques, e.g., SRRF and ESI, offering further imaging speed and quality enhancements in live-cell SR imaging (**Extended Data Fig. 9, Supplementary Video 3**, details in **Supplementary Note 5**).

### Rapid 4D live-cell SR imaging by Sparse-SACD

Rapid and intricate dynamic processes require fast SR imaging, and the temporal resolution improvement by SACD makes visualizing this activity feasible. Nevertheless, the real stable and high-fidelity live-cell SR imaging is always non-trivial, in which the available SNR is severely limited by photobleaching and phototoxicity. In this context, we introduce our previously developed Sparse deconvolution^25^ to maximize the utilization of accessible photon flux and split it into a pre-background-subtraction and a post-deconvolution constrained by using the continuity and sparsity penalties in *xy*-*t* (*z*) axes (**Methods, Fig. 4a, Supplementary Note 6**). Throughout 20-second consecutive SR imaging, we found the SNR in the raw recorded data decreased rapidly (**Fig. 4b, 4c, Supplementary Video 4**). Although being enhanced significantly against SOFI (**Fig. 4c**), such SNR reduction still induced a decrease in continuity and fidelity (**Fig. 4c**). Applying continuity and sparsity joint-constraint, the Sparse deconvolution-assisted SACD largely suppressed the potential artifacts, yielding high-fidelity continuous filaments (**Fig. 4c, Supplementary Fig. 4**).

**Fig. 4.**
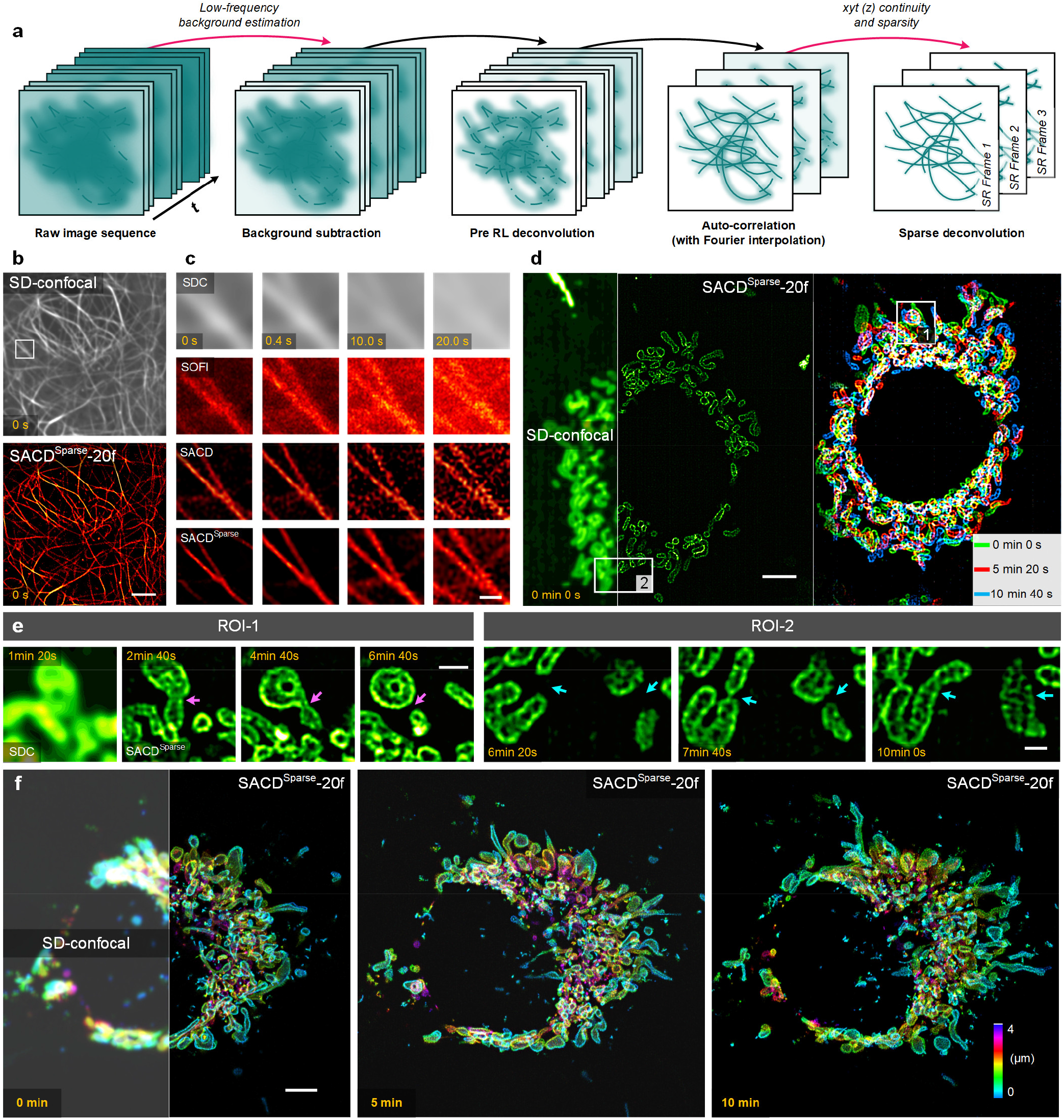
4D live-cell SR imaging using Sparse deconvolution-assisted SACD. (**a**) Workflow of Sparse-SACD. (**b**) A representative live COS-7 cell labeled with MAP4-Skylan-S imaged at 37°C by SD-confocal (top) and Sparse-SACD (‘SACD^Sparse^’) with 20 frames (bottom) (**Supplementary Video 4**). (**c**) From top to bottom: Representative montages of enlarged regions enclosed by the white box in (**b**) of SD-confocal (labeled as ‘SDC’), SOFI, SACD, Sparse-SACD with 20 frames. (**d**) From left to right: A representative live COS-7 cell labeled with Skylan-S-TOM20 imaged at 37°C imaged by SD-confocal (left) and Sparse-SACD (middle) at the first time-point, and the color-coded temporal projection of Sparse-SACD (right) (**Supplementary Video 5**). (**e**) Magnified montages of the corresponding white boxed regions in (**c**). The first one is the SD-confocal result (‘SDC’) and the others are Sparse-SACD results. (**f**) 4D imaging of a live COS-7 cell labeled with Skylan-S-TOM20 at 37°C using SD-confocal (left) and Sparse-SACD (right) (**Supplementary Video 6**). Scale bars: (**b, d, f**,) 5 μm; (**c, e**) 1 μm.

Next, we applied the Sparse-SACD to capture the live COS-7 cells (labeled with the Skylan-S-TOM20) over 10 minutes with a 40 seconds interval (**Fig. 4d, Extended Data Fig. 10, Supplementary Video 5**). The mitochondria were clearly resolved as hollow organelles, and their fine changes in morphology during fission (cyan arrows in **Fig. 4e**) and fusion (magenta arrows in **Fig. 4e**) were caught in several sequential SR recordings across minutes, which is generally acknowledged as a vital role in maintaining mitochondrial physiological function. On the other hand, under this live-cell condition, the previously modified SOFI methods exhibited severely aberrated SR images, especially at the end imaging stage (**Extended Data Fig. 7d-7e**, details in **Supplementary Note 7**). Beyond that, we also extended our SACD application to the challenging 4D SR imaging in whole live cells (**Fig. 4f, Supplementary Video 6**). A super-resolved 3D mitochondrial network in stretching was imaged across ten minutes. Compared to the traditional confocal system, the visibility of fine structure in depth has been improved thoroughly, and by use of sparsity-continuity constraint, the subtle whole-cell mitochondrial varying in time can be observed in high fidelity.

## DISCUSSION

>Abstractly, the individually fluctuating molecules distributed in the strong background are like the “*shining stars in the mist*”. Our core intention was to dispel such “*mist*” before the cumulant calculation, to enable high-quality SR under real physiological conditions. Using this mechanism, we have developed an easy-to-use algorithm, SACD, for efficient high-throughput and live-cell SR imaging. By fully leveraging the fluorescence fluctuation behavior from minimal datasets, our SACD significantly breaks the frame-number limitation in conventional SOFI, reducing the requisite frames by orders of magnitude. Utilizing our method, only 20 frames were employed instead of hundreds to thousands, offering the rapid imaging capability for insight into the biological structures at ∼100 nm scale. Then, our SACD was directly applied to a commercialized high-throughput SD-confocal microscopy system, achieving a stable ∼128 nm resolution over the entire centimeter-scale FOV. This demonstrated that SACD may be beneficial to high-throughput screening, for example, particle analyses^36^, or drug treatments^4^. Beyond that, it is also interesting to note that our framework is even feasible for analyzing a wide range of densities for single-molecule localization microscopy (SMLM) data^11^, allowing superior performance in high-density images (**Supplementary Fig. 5**).

On the other side, SIM is commonly applied to observe highly dynamic interactions in live cells. But its potential aberrations and distortions of the corresponding purpose-built illumination will lead to the fidelity deteriorating. These concerns highly compromise its expected usability in applications, especially its capacity for 3D imaging in thick and scattered samples^37, 38^. Another major crux of SIM is its unreliable robustness because the reconstruction process is excessively sensitive to noise conditions^35, 39^. These challenges potentially distort the real signals, even possibly leading to the failure of reconstruction, which may cause misinterpretations of biological analysis. The mechanism of our SACD not only effectively increases temporal resolving power, but also brings considerable benefits against SIM, such as liberating sophisticated optical design, being insensitive to noise conditions, and facilitating a large imaging field. Therefore, SACD may be another competent choice for fast and long-term live-cell SR imaging.

Empowering SR imaging with a real feasible spatiotemporal resolution for fluctuation-based approaches, the remaining possible expectation of SACD may focus on the further developments of diverse selectable protein tags with high on/off ratio for multicolor and *in vivo* imaging. Another expectation is that, when the necessary number of frames for calculating the higher-order cumulant exhibits insignificant increase, it may be considered to further extend the spatial resolutions and even facilitate quantitative studies, such as extraction of molecule density on cell membranes^40^. We have shown that the concept of SACD can directly achieve resolution improvement on a commercial system. Thus, we anticipate that the complete SACD toolkit will become a routinely used tool for biologists to dissect intricate and transient dynamics in live cells, and it may facilitate the microscopy-based screening with high-throughput, high-resolution, and low-cost that were previously inaccessible.

## METHODS

### SACD reconstruction

#### Fluctuations-based SR imaging

For simplicity, the sample to be imaged is usually regarded as composed of *N* single, independently fluctuating emitters^12^, located at position *r*_*k*_. These emitters are considered to have an individual time-dependent fluctuating molecular brightness, and the fluorescence signal at position *r* and time *t* can be given as:

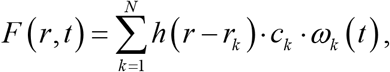

where the *h, c*, and *ω* represent the point-spread-function (PSF) of the corresponding microscope, the constant molecular brightness, and the time-dependent function of the molecular brightness fluctuating. As started by SOFI^12, 41^ which exploits such individual fluctuation character of each molecular, we can calculate the correlation cumulant for each pixel along the *t*-axial to enhance the resolution. Taking the second-order temporal cumulant *G*_2_ with zero time lag leads to

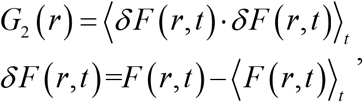

Where ⟨· ⟩_t_denotes the time averaging. Then if expanding the expression of *G*_2_, we can obtain the following formula.

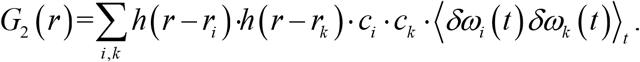

Assuming that each molecular is fluctuating individually (not correlating), the cross-correlation terms in this equation can be regarded as zero when *i* ≠ *k*. Therefore, such second-order temporal cumulant can be written as the simple sum of the squared PSF weighted by the corresponding constant brightness *γ*.

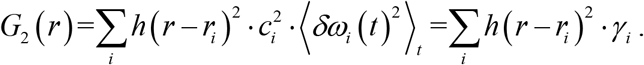

The width of the resulting PSF in this equation is reduced by a factor of 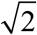, thus such second-order temporal cumulant can improve the resolution by ∼1.4 times at most.

#### Major steps of SACD reconstruction

However, if it is in real experiments, the real fluorescent content we imaged includes not only in-focus fluorescent molecules, but also out-of-focus and cytosol fluorescence background and baseline signals (**Extended Data Fig. 1a**), which may be not stationary equilibrium. The effective on/off contrast ratio is influenced by the unessential fluorescence, and the effective signal-to-noise ratio (SNR) condition dominated by inevitable readout noise, dark current noise, and shot noise of the sensor (**Extended Data Fig. 1a**), will affect the accuracy of cumulant calculation and the quality of the reconstrued image^19, 26, 33, 42, 43^. Thus, the real fluorescent content can be given by:

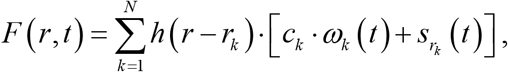

where the 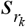 represents the mixed influence of the cytosol background and baseline noise signal at position *r*_*k*_. Then the *G*_2_ can be rewritten as:

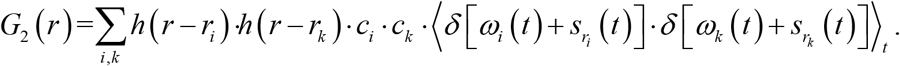

The cytosol background and baseline noise signal are usually assumed to be smoothly varying in the whole FOV, being spatiotemporally correlated. Thus, the cross-correlation terms in this equation cannot be treated as zero, which will compromise the ability of resolution enhancement. Such spatiotemporally correlated cytosol and baseline signals, as well as the noise components, can be reduced effectively by increasing the number of frames as demonstrated in the previous work^12^.

Here, we apply a pre-deconvolution step before the cumulant calculation as a tool to simultaneously reduce the noise, reject the out-of-focus and cytosol baseline signals, and improve the resolution. With this pre-deconvolution, the resolution enhancement ability of the temporal correlation cumulant can be used more efficiently. In specific, we employ the Richardson-Lucy (RL) deconvolution^28, 29^ without any regularizations as the first step of our SACD reconstruction. Using a few iterations, the RL deconvolution can filtrate the noise and sharpen the image without noticeable artifacts production. We used the accelerated RL deconvolution in this work:

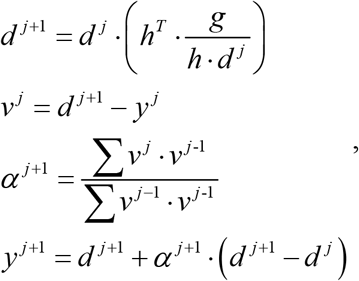

where *y* ^*n*+1^ is the image after *n*+1 iterations; *g* is the input image; and *h* is the PSF. The adaptive acceleration factor *α* was introduced by Andrews et al.^44^, representing the length of an iteration step, which can be estimated directly from experimental results. We charily stop its iteration at a very early stage (usually 10 iterations or the number automatically determined as in FRC-assisted SACD) to avoid destroying the independence of fluorescence fluctuation.

To match the improved spatial resolution by the subsequent auto-correlation cumulant calculation, we apply the Fourier interpolation^30^ to the image stack after the pre-deconvolution step, which can generate a smaller pixel size and larger pixel number, in the meanwhile keeping the real image content unchanged. After that, we calculate the auto-correlation cumulant of the RL processed, Fourier interpolated image sequence to improve the resolution. Then, considering that the resulting cumulant image is background-free, the deconvolution approach is demonstrated as a very suitable manner to improve the image resolution further^45^. At the final stage, a long step RL deconvolution is used on the single image after the auto-correlation reconstruction with the square of PSF, which further extends spatial resolutions along three dimensions. Ultimately, these four major steps included in the SACD reconstruction will achieve a more efficient twofold resolution enhancement, resulting in the massive reduction of the necessary sequence length.

### FRC-assisted parameterless SACD

To make our SACD method easy to use, we apply the Fourier ring correlation (FRC) to determine the optimal iteration times automatically (**Extended Data Fig. 1b**), which has been used as an effective resolution criterion for SR and electron microscopy^21, 22, 46^. FRC calculation needs statistically independent image pairs, that share identical details (correlated), but different noise realizations (uncorrelated), which describes the highest acceptable frequency component of distance between two signals. Here we use the FRC to estimate the reliable cutoff frequency to determine the best iteration automatically. In another word, we intend to obtain a higher reliable cutoff frequency, which indicates better image quality or resolution.

In specific, we split the raw image sequence in half (odd and even frames) and subsequently these subsets were averaged to generate the required two frames for the FRC calculation. Then, we estimate the FRC value of each iteration result, and in practice it is usually a quadric curve along the iteration times. The iteration of the pre-RL-deconvolution should be stopped before influencing the auto-correlation calculation (**Extended Data Fig. 3**). Therefore, we stop the pre-RL-deconvolution at half iteration times of maximum resolution (lowest FRC value). On the other hand, when considering that the resulting auto-correlation image is background-free, the long step of RL deconvolution will be tending to offer superior and stationary performance. Thus, we stop the post-RL-deconvolution when reaching the maximum resolution to maximize the resolution improvement (**Extended Data Fig. 1b**).

In addition to the determination of the optimal iteration times, we can also utilize FRC to estimate the PSFs of the corresponding images (**Extended Data Fig. 1b**). In previous work, the global FRC value was used to estimate the PSF of the image, and such estimated PSF was applied in the blind RL deconvolution^22^. However, it is notable that the FRC estimates the effective PSF (the reliable frequency components), rather than the native resolution of the microscope system. The large noise amplitude will dominate the frequency components even inside the optical transfer function (OTF), which will influence the FRC to infer a larger resolution. On the other hand, the RL deconvolution requires the native resolution of the corresponding microscope system. Thus, this FRC-based PSF estimation introduces the assumption that the corresponding image is under sufficiently high SNR conditions, which is usually unsatisfied for the raw images. Therefore, we calculate the PSF with the Born-Wolf model^47^ in the pre-deconvolution step, and execute the PSF estimation using FRC on the noise-free auto-correlation reconstruction result in the post-deconvolution step.

### Sparse-SACD reconstruction

Although the pre-RL-deconvolution has been used in SACD to reduce the noise and background, it still has limits on dealing with the images containing large noise and strong background signals. For sensitive live-cell SR imaging, the achievable signal-to-noise ratio (SNR) is severely limited by photobleaching and phototoxicity and it will decrease continually during the long-term image sequence acquisition. Predictably, the SACD reconstructed result will suffer from artifacts when the SNR is lower to a certain extent. This issue could be resolved by increasing the number of captured images. Besides, when the image contains a strong cytosol signal, sometimes even increasing the sequence length still cannot fully remove such background. Remarkably, these two problems exist commonly in fast and long-term live-cell imaging.

To maintain the temporal resolution of SACD, we involved the previously developed Sparse deconvolution^25^ in our SACD reconstruction (Sparse-SACD) to address the above limitations. The Sparse deconvolution is a constrained deconvolution algorithm that can reduce the artifacts and increase the spatial resolution. The framework contains two main steps, background subtraction and iterative deconvolution with sparsity and continuity constraints. In specific, we use background subtraction on the raw images to remove the unnecessary low-frequency baseline signal and strong cytosol background (optional, before **Step 1** in SACD). The constrained iterative deconvolution is applied at the last step (replacing **Step 4** in SACD), which enables the resolution improvement further while maintaining the image quality (minimizing the artifacts).

We use an iterative wavelet decomposition method to estimate and subtract the background. The background is iteratively estimated from the lowest frequency wavelet bands of the images, and in each iteration, all values of the image above the current estimated background level are clipped. The constrained iterative deconvolution is given by:

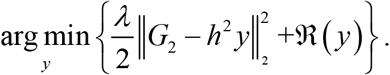

The first term on the left side of the equation is the fidelity term, representing the distance between the recovered image *y* and the result obtained with the calculation of auto-correlation image *G*_2_. *h* is the PSF of the optical system. The second term represents the continuity and sparsity constraints, which is given by:

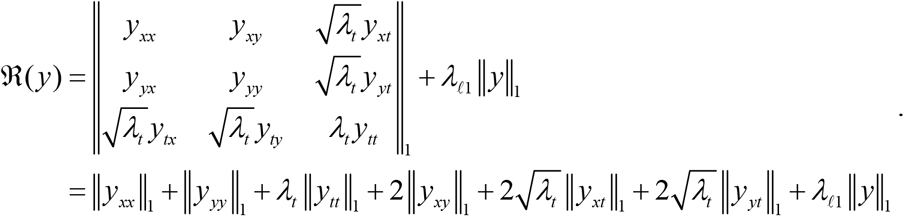

The first term is the continuity prior regarding the structural continuity along the *xy*-*t*(*z*) axes, and the second term is the sparsity constraint. *λ*_*t*_ denotes the regularization parameters that present the continuity along the *t*-axis, which can be turned off upon imaging objects moving at high speed. The subscript *xx* is the two-order derivation operator in the *x*-direction. *λ* and 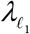 denote weight factors to balance image fidelity with sparsity.

The high-frequency artifacts caused by the readout noise dark current and shot noise are mostly non-continuous along the *xy*-*t*(*z*) axes. Moreover, to ensure sufficient Nyquist sampling criteria for maximal spatial resolution dictated by the optics, the PSF must occupy more than 3 × 3 pixels in space, which constitutes the basis for the continuity along *x* and *y* axes. On the other hand, fluorescence microscopy achieves high resolution and contrast by labeling designated structures, which are always sparse in the spatial domain in terms of the existence of a fraction of nonzero value. Moreover, the fluorescence signal is blurred by the optical system, and it is accompanied by noise when captured by the photoelectric detector. Therefore, the real signal is always more sparse than the recorded signal^25^. Therefore, the continuity prior and the sparsity prior are involved in our Sparse-SACD to constrain the final step deconvolution for reducing the artifacts and improving the resolution. As described previously, to solve such convex optimization problem, we split it into constraint reconstruction and deconvolution subproblems. We solve these two subproblems using a two-step operation, to gradually and effectively search for the final solution *y*. In specific, the Split-Bregman framework^48^ is employed to execute the reconstruction of continuity and sparsity weighted constraints and we use RL deconvolution for the iterative deconvolution subproblem.

### SD-confocal setup

A commercial SD-confocal microscope system, i.e., Spin-disk system (Dragonfly, Andor) based on an inverted fluorescence microscope (DMi8, Leica) with an oil immersion objective (100×/1.3 oil, Plan Apo, Leica), is used in this work. Four laser beams of 405 nm, 488 nm, 561 nm, and 647 nm were combined with the SD-confocal microscope. The images were captured either by a sCMOS camera (Zyla 4.2 Plus, Andor) or an EMCCD camera (iXon Ultra 888, Andor).

### SIM setup

The SIM system is based on a commercial inverted fluorescence microscope (IX83, Olympus) equipped with an objective (100×/1.7 HI oil, APON, Olympus) and a multiband dichroic mirror (DM, ZT405/488/561/640-phase R; Chroma) as described previously^35^. In short, laser light with wavelengths of 488 nm (Sapphire 488LP-200) and 561 nm (Sapphire 561LP-200, Coherent) and acoustic optical tunable filters (AOTF, AA Opto-Electronic, France) were used to combine, switch, and adjust the illumination power of the lasers. A collimating lens (focal length: 10 mm, Lightpath) was used to couple the lasers to a polarization-maintaining single-mode fiber (QPMJ-3AF3S, Oz Optics). The output lasers were then collimated by an objective lens (CFI Plan Apochromat Lambda 2× NA 0.10, Nikon) and diffracted by the pure phase grating that consisted of a polarizing beam splitter, a half-wave plate, and the SLM (3DM-SXGA, ForthDD). The diffraction beams were then focused by another achromatic lens (AC508-250, Thorlabs) onto the intermediate pupil plane, where a carefully designed stop mask was placed to block the zero-order beam and other stray light and to permit passage of ±1 ordered beam pairs only. To maximally modulate the illumination pattern while eliminating the switching time between different excitation polarizations, a home-made polarization rotator was placed after the stop mask^35^. Next, the light passed through another lens (AC254-125, Thorlabs) and a tube lens (ITL200, Thorlabs) to focus on the back focal plane of the objective lens, which interfered with the image plane after passing through the objective lens. Emitted fluorescence collected by the same objective passed through a dichroic mirror, an emission filter, and another tube lens. Finally, the emitted fluorescence was split by an image splitter (W-VIEW GEMINI, Hamamatsu, Japan) before being captured by a sCMOS (Flash 4.0 V3, Hamamatsu, Japan) camera.

### Wide-field and TIRF microscopy setup

The three phases of structured illumination under the same orientation can be averaged to a uniform wide-field illumination. Taking advantage of that, we use the SIM setup described in the above section to generate the wide-field images by integrating three frames (corresponding to three phases of structured illumination) on the camera plane, which enables more flexible cross-validation of SIM and SACD results. In another word, we employ the identical commercial inverted fluorescence microscope (IX83, Olympus) equipped with an objective (100×/1.7 HI oil, APON, Olympus)and a sCMOS (Flash 4.0 V3, Hamamatsu, Japan) camera to capture the wide-field images for our SACD reconstruction.

### High-throughput SR image acquisition and reconstruction

For high-throughput imaging of microtubules in COS-7 cells, we used a commercialized solution, i.e., the same type SD-confocal microscope equipped with a montage stitching tool (Dragonfly, Andor), to define the positions of 704 field-of-view (FOVs) (each contained 1024 × 1024 pixels) of 2.0 mm × 1.4 mm on a 32 × 22 grid, with 5% overlap. The stage was automatically shifted to each of the 704 positions. The exposure time at each FOV was ∼30 ms (∼40 ms including readout time), and the time series is 20 frames at each position. The total imaging time is ∼9 minutes 23 seconds, and the stage moving time is ∼10 minutes, so the total acquisition time is ∼20 minutes.

Then the full stitched image stack (30193 × 20913 × 20 pixels) was cut to 759 (33 × 23) partly overlapping (12.5%) tiles, 1024 × 1024 × 20 pixels each. We employed SACD reconstruction on these tiles individually, and the resulting SR images were applied with a histogram matching (previous tile as reference) to normalize the intensity for a better subsequent fusion quality. Finally, the tiles were stitched back to a full SR image (60386 × 41826), and this resulting image was downsampled by 12 times (5032 × 3486 pixels) for the visualizations in **Fig. 3a** and **Supplementary Fig. 3a**.

### Cell maintenance and preparation

COS-7 cells were cultured in high-glucose DMEM (GIBCO, 21063029) supplemented with 10% fetal bovine serum (FBS) (GIBCO) and 1% 100 mM sodium pyruvate solution (Sigma-Aldrich, S8636) in an incubator at 37°C with 5% CO^2^ until ∼75% confluency was reached. MCF-7 cells were cultured in MEM (Thermo fisher, 11095072) supplemented with 10% fetal bovine serum (FBS), 0.01 mg/ml human recombinant insulin (Sigma, I9278) and 1% 100 mM sodium pyruvate solution. For the SD-confocal imaging experiments, 35-mm glass-bottomed dishes (Cellvis, D35-14-1-N) were used. For the wide-field and 2D-SIM imaging experiments, cells were seeded onto coverslips (H-LAF 10L glass, reflection index: 1.788, diameter: 26 mm, thickness: 0.15 mm, customized) coated with 0.01% Poly-L-lysine solution (Sigma, P4707) for 10 minutes and washed twice with Sterile Water before seeding transfected cells.

### Immunofluorescence

The COS-7 cells were grown in 35-mm glass-bottomed dishes overnight and rinsed with PBS, then immediately fixed with prewarmed 4% PFA (Santa Cruz Biotechnology, sc-281692) for 10 mins. After three washes with PBS, cells were permeabilized with 0.1% Triton® X-100 (Sigma-Aldrich, X-100) in PBS for 10 mins. Cells were blocked in 5% BSA/PBS for 1h at room temperature. Mouse anti-Tubulin DM1a (Sigma, T6199) was diluted to 1:100 and stained cells in 2.5% BSA/PBS blocking solution for 2 h at room temperature. The cells were then washed with PBS five times for 10 mins per wash and stained with Biotin-XX goat anti-mouse IgG antibody (Invitrogen, B2763) at room temperature in PBS with 1:200 for 1 h. The cells were then washed with PBS five times for 10 mins per wash and stained with Qdot™ 525 Streptavidin Conjugate (Invitrogen, Q10143MP) or Qdot™ 605 Streptavidin Conjugate (Invitrogen, Q10103MP) at room temperature in PBS with 1:50 for 1 h. Finally, cells were washed five times with PBS and imaged. In addition, according to the Qdot property, all kinds of Qdot were excited at 405 nm, and the number of Qdot™ 525 and Qdot™ 605 means emission wavelength.

### Live-cell samples

To label the mitochondria/microtubules in live cells, COS-7 cells were transfected with Skylan-S-TOM20 and MAP4-Skylan-S^19^. The transfections were executed using LipofectamineTM 2000 (Thermo Fisher Scientific, 11668019) according to the manufacturer’s instructions. After transfection, cells were plated in glass-bottomed dishes. The Skylan-S was under sequential illumination with a 405 nm laser (low power) when imaging. In addition, live cells were imaged in a complete cell culture medium containing no phenol red in a 37°C live cell imaging system.

### Live-cell SRRF dataset

The GFP-tagged microtubules in live HeLa cells (**Supplementary Fig. 8**) were imaged in TIRF mode with a TIRF objective (100×/1.46 NA, Oil, Plan Apochromat, Zeiss) and an additional 1.6× magnification with 488-nm laser illumination^13^.

### Simulation protocol of the fluctuating fluorescence samples

Ground truth structures were created on a fine grid (10 nm or 20 nm). Then, following simulation procedures in the SOFI simulation tool^49^, the blinking behavior of each emitter is modeled as a time-continuous Markovian process. Using that, we generated temporal blinking sequences of the two fluorescence emitters and convolved these image sequences with a 220 nm PSF subsequently. The resulting images were then subsampled multiple times for a pixel size of 60 nm. For the ‘Non-bias’ condition in **Extended Data Fig. 4a**, only the Poisson noise was involved in the images. To synthesize more realistic experimental imaging conditions, the cytosol background, out-of-focus light, Gaussian readout noise, and baseline background under different levels (‘low’, ‘medium’, and ‘high’ in **Extended Data Fig. 4b-4d, Extended Data Fig. 5**, and **Supplementary Fig. 6**; ‘ultralow’ in **Supplementary Fig. 9**) were sequentially included in the images.

### Performance metrics

#### FRC resolution

The calculation of FRC resolution requires two independent frames of identical contents under the same imaging conditions^21^. For the case of SOFI and SACD, these two frames were generated by splitting the raw image sequence into two image subsets (the first 20 frames and the last 20 frames) and reconstructing them independently. Regarding the case of wide-field and SIM (**Fig. 1e**), the raw image sequence was divided into two image subsets (the first 20 frames and the last 20 frames), and subsequently these subsets were averaged to create the required two frames.

#### Structural Similarity (SSIM)

The SSIM^34^ is defined as:

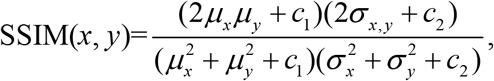

where *x* represents the reference which is ground-truth structures convolved with a 110 nm size PSF, and *y* denotes the corresponding SR reconstructions. The *μ*_*x*_ and *μ*_*y*_ are the averages of *x, y*; *σ*_*x, y*_ is the covariance of *x* and *y*; *σ*^2^are the variances; and *c*_1_, *c*_2_ are the variables used to stabilize the division with a small denominator.

#### Ring quality ratio (RQR)

Because one two-peak analysis may be not sufficient to validate that the ring is fully resolved^25^, to quantitatively evaluate the reconstruction quality, we divided two circular/annular areas based on the corresponding ground-truth diameters. Following the modulation transfer function, we defined the RQR calculation:

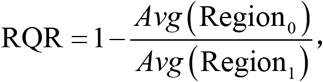

where *Avg* calculates the mean intensity of pixels within the corresponding area. The Region_0_ and Region_1_ represent the double annulus region (in red, **Extended Data Fig. 5h**) and the region in between (in yellow, **Extended Data Fig. 5h**). To avoid overconfident determination, we used a threshold of 0.4 to determine whether the ring structures are successfully resolved.

#### Standard deviation (STD)

To measure the reconstruction uncertainty (STD of reconstruction) of fluctuation-based SR methods, following ref^18^, the Jackknife resampling was used to create a reconstruction stack. Then, the resulting image stack was performed the STD calculation:

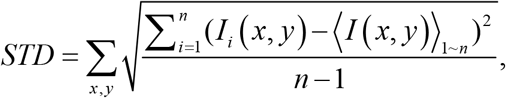

where *n* is the sequence length; (*x, y*) denotes the two-dimensional coordinate; *I*_*i*_ (*x, y*) represents the intensity of the *i-*th image in the sequence; and < *I* (*x, y*) >_1∼*n*_ is the averaged intensity of the sequence.

### Reconstruction algorithms and their parameters for fluorescence fluctuation-based data

We compare performances against three other modified SOFI methods: SOFI-wavelet^32^, SOFI-RL^31, 45^, and RD-Covar^33^ (covariance calculation followed by RL deconvolution); and 5 other freely available software packages: SRRF^13^, ESI^15^, HAWK^50^, ThunderSTORM^51^, and bSOFI^45^. Notably, because the three modified SOFI methods are not open-sourced, we implemented them with adherence to the published work. We will describe how we configured each of these methods in turn.

#### SOFI-wavelet

A 2D Fourier upsampling operation was performed on the raw stack, followed by a two-step wavelet-based filter. The first step is equal to a high-pass filter along the *t*-axial for each pixel to remove background fluctuations, in which we used 2D Daubechies-4 wavelet filters to decompose the signal up to the 7th level and remove the lowest component. The second step is similar to a low-pass filter on each image to moderate readout noise, in which we used 2D Daubechies-4 wavelet filters to decompose the signal up to the 4th level and cut off the highest component. After that, the 2^nd^-order cumulant was employed on the image stack.

#### SOFI-RL

A 2D Fourier upsampling operation was performed on the raw stack, followed by a 2^nd^-order cumulant calculation. Then we applied the RL deconvolution to the resulting image.

#### RD-Covar

A 2D Fourier upsampling operation was performed on the raw stack, followed by a 2^nd^-order covariance calculation, and then we applied the RL deconvolution to the resulting image. The covariance calculation is defined as:

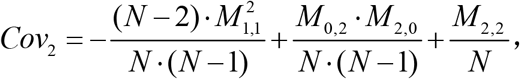

where *M*_*m,n*_ = ⟨*δ F*^*m*^ (***r***, *t*) ·*δ F*^*n*^ (***r***, *t* +*τ*) ⟩; *δ F* (***r***, *t*) = *F* (***r***, *t*) −⟨*F* (***r***, *t*) ⟩; *F* (**r**, *t*) represents the recorded signal intensity at position ***r*** and time *t*; and *N* is the sequence length. We used *τ* as 0 in this work.

#### SRRF ImageJ plugin

Images labeled with ‘SRRF’ were reconstructed by the temporal radiality average method, with default parameters, except the ‘Radiality magnification’ as ‘4’ for **Fig. 1c** and ‘4’ for **Supplementary Fig. 8**. Images labeled with ‘SRRF-TRC’ and ‘SRRF-TRAC’ in **Extended Data Fig. 9** were reconstructed by the temporal radiality average and the temporal radiality auto-correlation methods respectively, with default parameters, except the ‘Radiality magnification’ as ‘2’. *ESI ImageJ plugin*. Images labeled with ‘ESI-2nd’ in **Fig. 1c** and **Extended Data Fig. 9** and ‘ESI’ in **Supplementary Fig. 5** were reconstructed by 2^nd^ order and 4^th^ order ESI respectively, with default parameters.

#### HAWK ImageJ plugin

The image labeled with ‘HAWK-SOFI’ in **Extended Data Fig. 9** was reconstructed by HAWK pre-processing with default parameters followed by SOFI.

#### ThunderSTORM ImageJ plugin

The image labeled with ‘ThunderSTORM’ in **Supplementary Fig. 5** was reconstructed by the maximum likelihood fitting with the integrated PSF method and multi-emitter fitting enabled. Drift correction was performed post-localization and images were rendered using an average shifted histogram approach.

#### bSOFI MATLAB library

The image labeled with ‘bSOFI in **Supplementary Fig. 5** was reconstructed by the 4^th^ order bSOFI with default parameters.

### Image rendering and processing

#### Diffraction-limit image rendering

We applied a rolling ball background subtraction on the diffraction-limit image and filtered the resulting image with a small Gaussian kernel (1 pixel in radius) for removing readout noise.

#### SOFI image rendering

We followed the bSOFI^45^ configuration to visualize the SOFI images. Because the intensity after 2^nd^ order cumulant will be scaled to its squared value, we took the square root on the image to linearize the brightness. To reduce the amplified noise induced by this gamma correction, small values (typically 2% of the maximum value) are truncated, and subsequently the image is re-convolved with a small Gaussian kernel (FWHM as the corresponding doubled resolution). Notably, to avoid canceling the resolution improvement of the cumulant, the SOFI images without post-deconvolution were directly visualized by its raw version with brightness adjustment.

#### Colormap and software

The color map ‘biop-12colors’ was used to color code the 3D volumes in **Fig. 2e** and **Fig. 4f**. The color map ‘Isolum’^52^ was used to color code the 3D volume in **Fig. 2b**. The color map ‘SQUIRREL-FRC’^53^ was used to color code the FRC maps in **Fig. 3a** and **Supplementary Fig. 3e**. The color map ‘Morgenstemning’^52^ was applied to show the Fourier transform results in **Extended Data Fig. 6c**. The color map ‘SQUIRREL-errors’^53^ was used to color code the resolution enhancing map in **Supplementary Fig. 3e**. The color map ‘Jet’ was used to show the time series in **Supplementary Fig. 8h**, and the intensity amplitudes in **Extended Data Fig. 10b**. All data processing was achieved using MATLAB and ImageJ. All figures were prepared with MATLAB, ImageJ, Microsoft Visio, and OriginPro, and videos were all produced with our light-weight MATLAB framework, which is available at https://github.com/WeisongZhao/img2vid.

## Data availability

All the data that support the findings of this study are available from the corresponding author on reasonable request.

## Code availability

The SACD in MATLAB library can be found at https://github.com/WeisongZhao/SACDm, and its corresponding ImageJ plugin can be found at https://github.com/WeisongZhao/SACDj.

## Acknowledgments

H. L. acknowledges support by the State Key Laboratory of Robotics and Systems. S. Z. acknowledges support by the Boya Postdoctoral Fellowship of Peking University. L. C. acknowledges support by the High-performance Computing Platform of Peking University. H. L. acknowledges support by grants from the National Natural Science Foundation of China (61805057); the Young Elite Scientists Sponsorship Program (2018QNRC001); and the Natural Science Foundation of Heilongjiang Province (YQ2021F013). L. C. acknowledges support by grants from the National Natural Science Foundation of China (92054301, 81925022, 31821091, 91750203), the National Science and Technology Major Project Program (2016YFA0500400); and the Beijing Natural Science Foundation (Z20J00059).

## Author contributions

H. L., L. C., and W. Z. supervised the project; W. Z. and H. L. initiated and conceived the research; W. Z. developed the algorithm and implemented the corresponding software; W. Z. and S. Z. designed the experiments; S. Z. performed the experiments and collected the data; W. Z. analyzed the data and prepared the figures with the contribution of X. D.; W. Z. performed the simulations with the contribution of Z. H., X. D., L. Q., X. L., and X. W..; Z. H. and W. Z. prepared the videos; H. M., Y. J., Y. H., Xumin D., G. H., C. G., and J. T. participated in discussions during the development of the manuscript; W. Z., H. L., and L. C. wrote the manuscript with input from all authors; All authors participated in the discussions and data interpretation.

## EXTENDED DATA FIGURES

**Extended Data Fig. 1.**
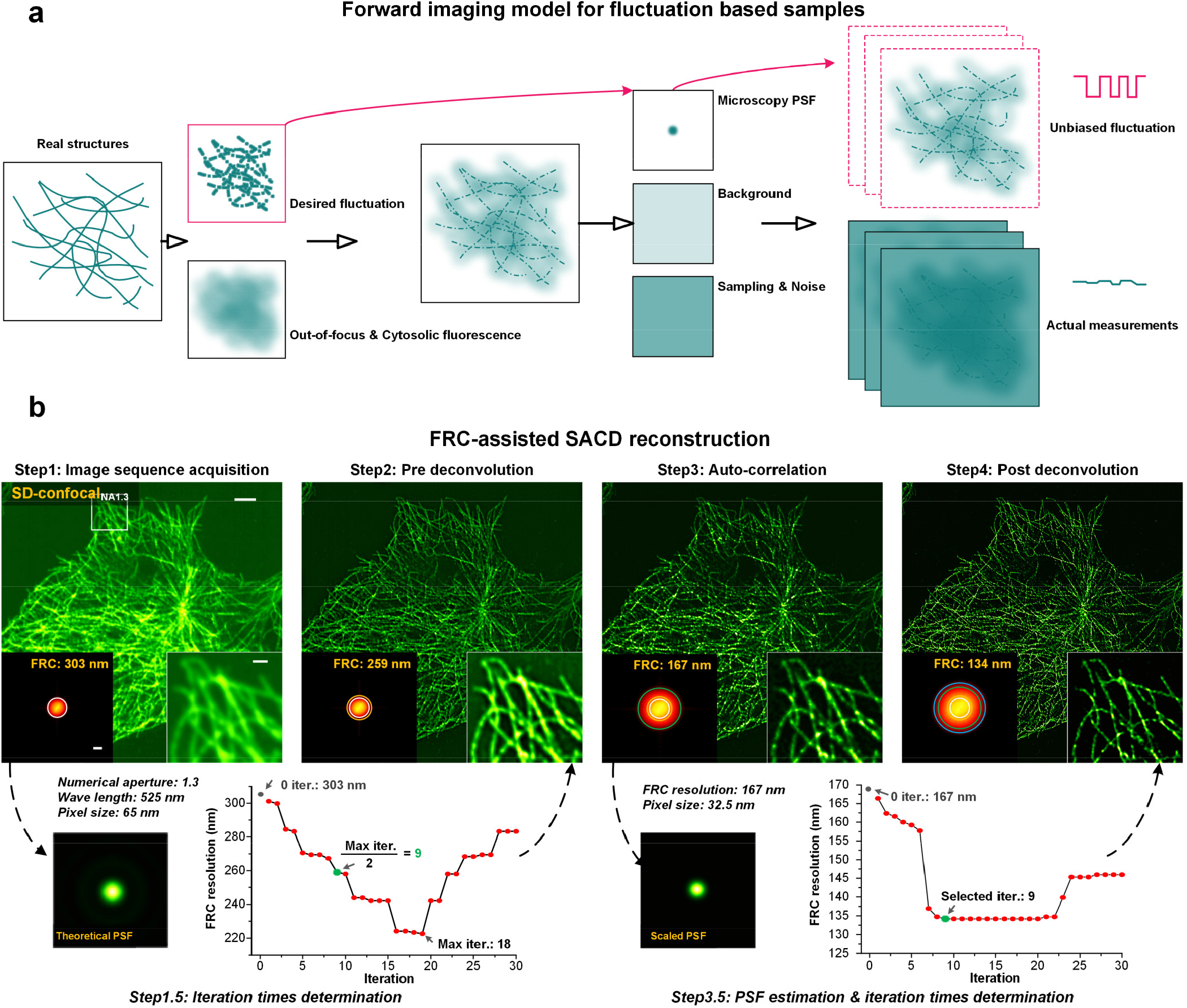
The forward imaging model of fluctuation-based samples and reconstruction workflow of the FRC-assisted parameterless SACD. (**a**) If it is in real experiments, the real fluorescent content we imaged includes not only in-focus fluorescent molecules (‘Desired fluctuation’), but also out-of-focus and cytosol fluorescence background. Then the fluorescence signal accompanied with baseline signal (‘Background’) is encoded by the microscopy PSF, and sampled by the sensor (‘Sampling & Noise’). The actual measurements will always deviate from the desired fluctuation model from SOFI (‘Unbiased fluctuation’, see also **Methods**). (**b**) Workflow and representative example (a COS-7 cell labeled with QD525) of the FRC-assisted parameterless SACD. Left insets of top panel: Fourier transforms, and the corresponding FRC resolutions of different stages are labeled on the top right; Right insets of top panel: Magnified views from the white box. In **Step1**, the image sequence for reconstruction is captured. In **Step 1.5**, the PSF of raw data for the pre-deconvolution is calculated by the theoretical model. In **Step2**, we perform RL deconvolution for each image and it will be stopped at half of the iteration of the maximal resolution (lowest FRC value). In **Step3**, we calculate 2^nd^ order auto-correlation cumulant. Then in **Step 3.5**, we estimate the FRC resolution of the resulting image to calculate PSF for post-deconvolution. In **Step4**, we employ post RL deconvolution for the auto-correlation image and it will be stopped when reaching the maximum resolution. Scale bars: (**b**, main panel and left inset) 5 μm; (**b**, right inset) 1 μm.

**Extended Data Fig. 2.**
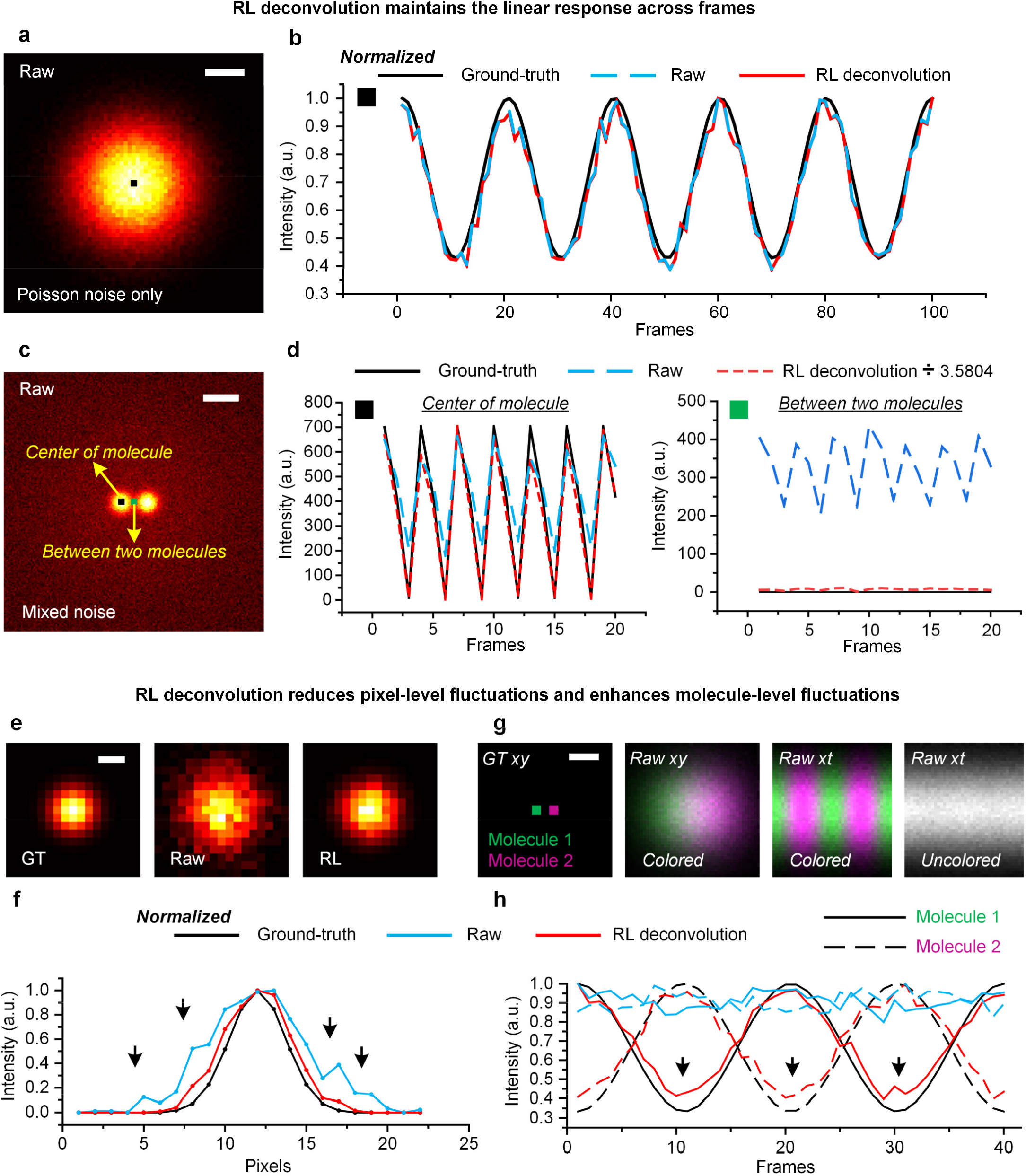
Characterizing properties of the pre-RL deconvolution. (**a**) A representative image of a single molecule with a cosine intensity fluctuation convolved by a 220 nm Gaussian PSF before adding Poisson noise only. (**b**) Normalized intensity profiles of ground-truth (black), raw (blue), and RL deconvolution (red) for the temporal response indicated by the black point in (**a**). (**c**) A representative image of two blinking molecules convolved by a 220 nm Gaussian PSF before adding Poisson noise, ∼2% Gaussian noise, and background noise. (**d**) Intensity profiles of ground-truth (black), raw (blue), and RL deconvolution (red, divided by 3.5804) for the temporal response indicated by the black point (left) and green point (right) in (**c**). (**e**) A single molecule convolved by a 155 nm Gaussian PSF as ground-truth (‘GT’; left), and convolved by a 220 nm Gaussian PSF before adding Poisson noise (‘Raw’; middle) followed by RL deconvolution (‘RL’; right). (**f**) Intensity profiles of ground-truth (black), raw (blue), and RL deconvolution (red). Black arrows indicate the pixel-level fluctuations in the raw capture (blue). (**g**) Representative images of two molecules with opposite cosine intensity fluctuations convolved by a 220 nm Gaussian PSF before adding Poisson noise, ∼2% Gaussian noise, and background noise. From left to right: Ground-truth molecules in *xy*; Colored rapture in *xy*; Colored raw capture in *xt*; Uncolored raw capture in *xy*. (**d**) Normalized intensity profiles of ground-truth (black), raw (blue), and RL deconvolution (red) for the temporal response indicated by the magenta point (dashed lines) and green point (solid lines) in (**g**, 1^st^ column). Details in **Supplementary Note 1**. Scale bars: (**a, e, g**) 100 nm; (**c**) 500 nm.

**Extended Data Fig. 3.**
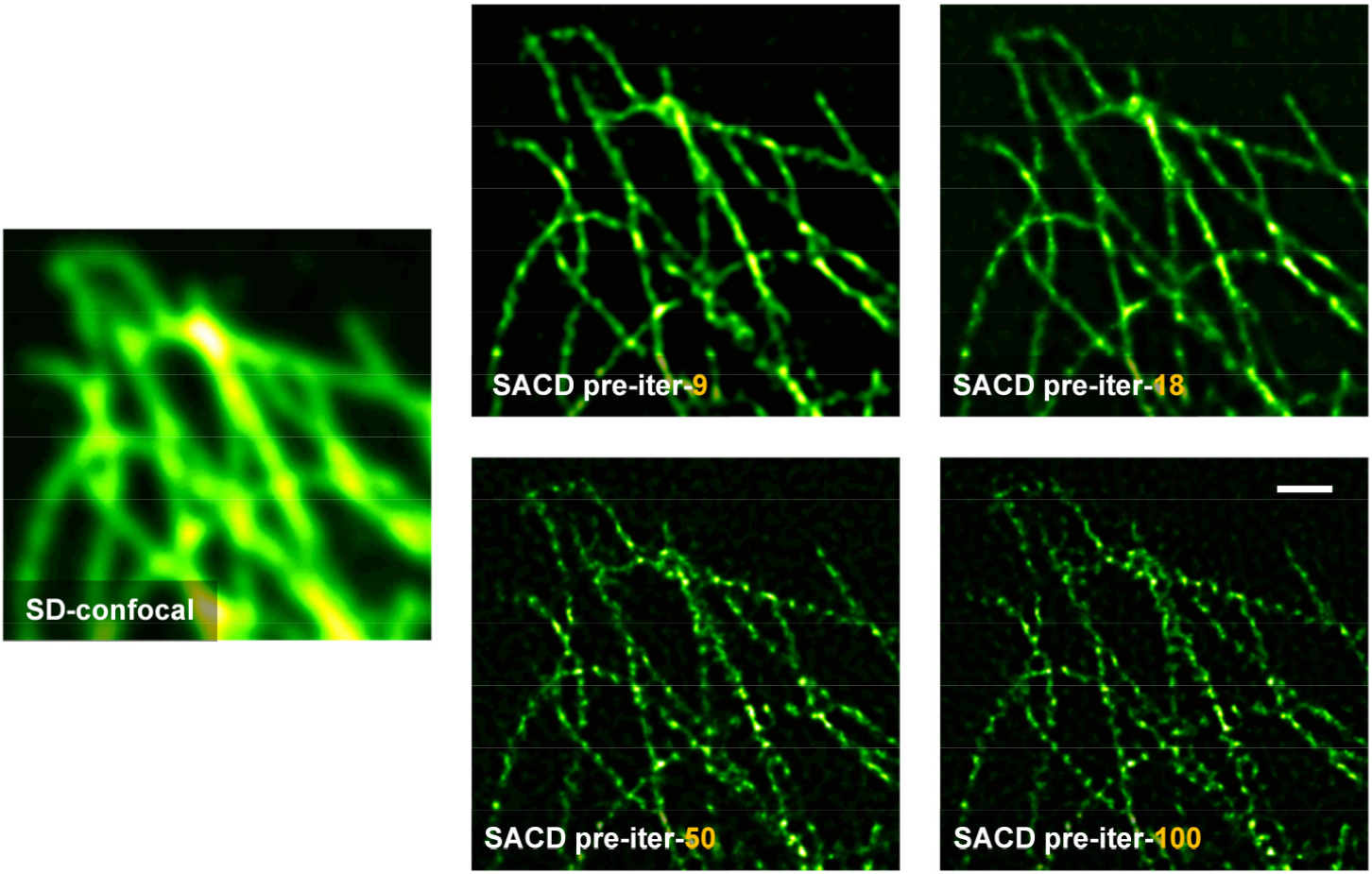
Comparisons of the pre-deconvolution using different iteration times. Microtubule filaments in a COS-7 cell labeled with QD_525_ (*c*.*f*., **Extended Data Fig. 1b**) imaged by SD-confocal (left, averaged by 20 frames) and SACD under 9 (top middle), 20(top right), 50 (bottom middle), and 100 (bottom right) iteration times of pre deconvolution configurations. Scale bar: 1 μm.

**Extended Data Fig. 4.**
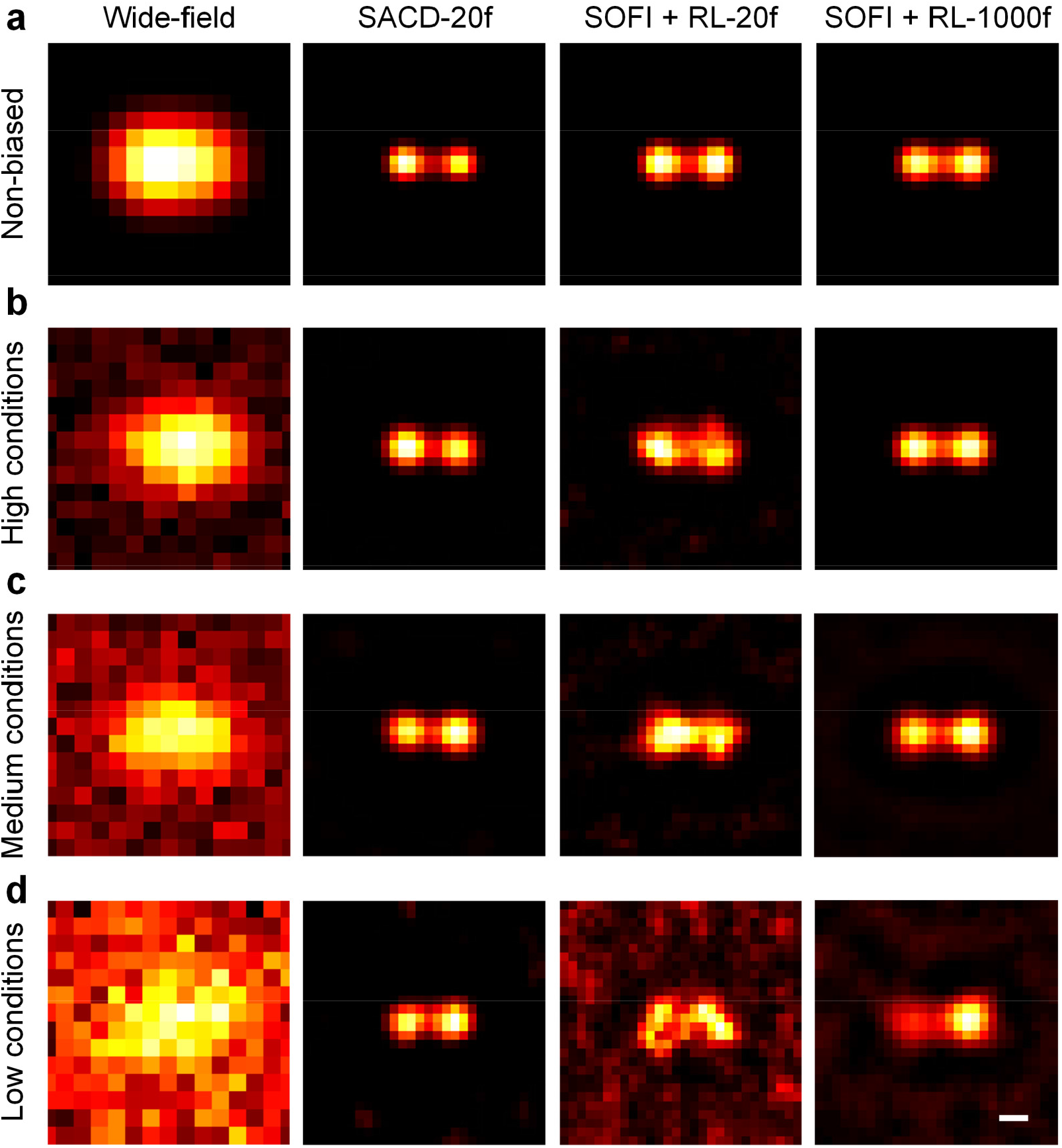
Simulation validation of SACD and SOFI reconstructions under different imaging conditions. (**a-d**) From left to right: Wide-field image, 20-frame SACD reconstruction, 20-frame SOFI reconstruction followed by RL deconvolution, and 1000-frame SOFI reconstruction followed by RL deconvolution. Two molecules 160 nm apart were created at a 10 nm size pixel grid. We generated temporal blinking sequences of the fluorescence emitters (**Methods**), and convolved them with a 220 nm PSF subsequently. The resulting image stack was downsampled 6 times (pixel size 60 nm). The ‘Non-biased’ condition (**b**) represents that only Poisson noise was considered in the imaging process. The high (**c**), medium (**d**), and low (**e**) conditions denote the corresponding high, medium, and low SNRs and effective on/off contrast ratios (**Methods**). These conditions involved not only Poisson noise and Gaussian readout noise, but also the smooth fluctuating cytosol and out-of-focus background. Scale bar: 100 nm.

**Extended Data Fig. 5.**
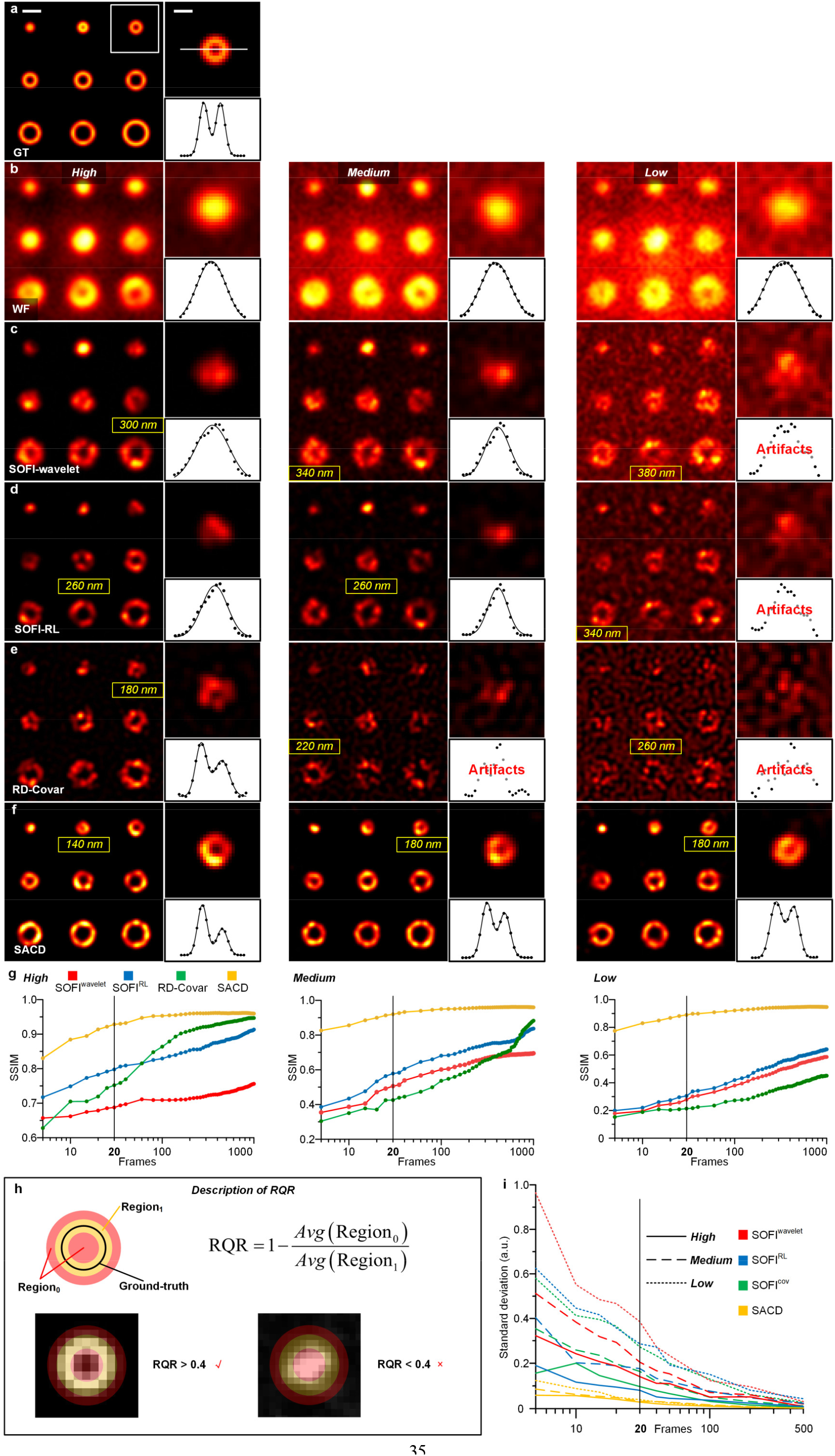
Simulation comparisons of different SOFI methods and SACD under different SNRs using ring structures. (**a**) The synthetic ring structures were convolved with a 110 nm PSF and down-sampled 3 times (pixel size 30 nm) as ground truth (Methods). (**b-f**) Wide-field (WF, **b**) raw images and SOFI-wavelet (**c**), SOFI-RL (**d**), RD-Covar (**e**), and SACD (**f**) reconstructions using 20-frame under ‘*High*’ (right), ‘*Medium*’ (middle), and ‘*Low*’ (left) SNR conditions. The structures were convolved with a 220 nm PSF, down-sampled 6 times before adding the cytosol background, out-of-focus light, Poisson noise, Gaussian readout noise, and baseline background under three different levels to be the raw WF image stacks. Yellow numbers represent the minimum resolvable diameters of ring structures according to the RQR criterion. The sub-images at the upper right corner are the enlarged view from white box. The sub-images at the lower right corner are the intensity profiles and multiple Gaussian fitting from white line. (**g**) The SSIM curves of different methods under ‘*High*’ (right), ‘*Medium*’ (middle), and ‘*Low*’ (left) SNR conditions. (**h**) The calculation method of ring quality ratio (RQR) criterion. (**i**) The STD curves of different methods under SNR conditions. Details in **Supplementary Note 2**. Scale bars: (**a**) 300 nm; (**a**, inset) 150 nm.

**Extended Data Fig. 6.**
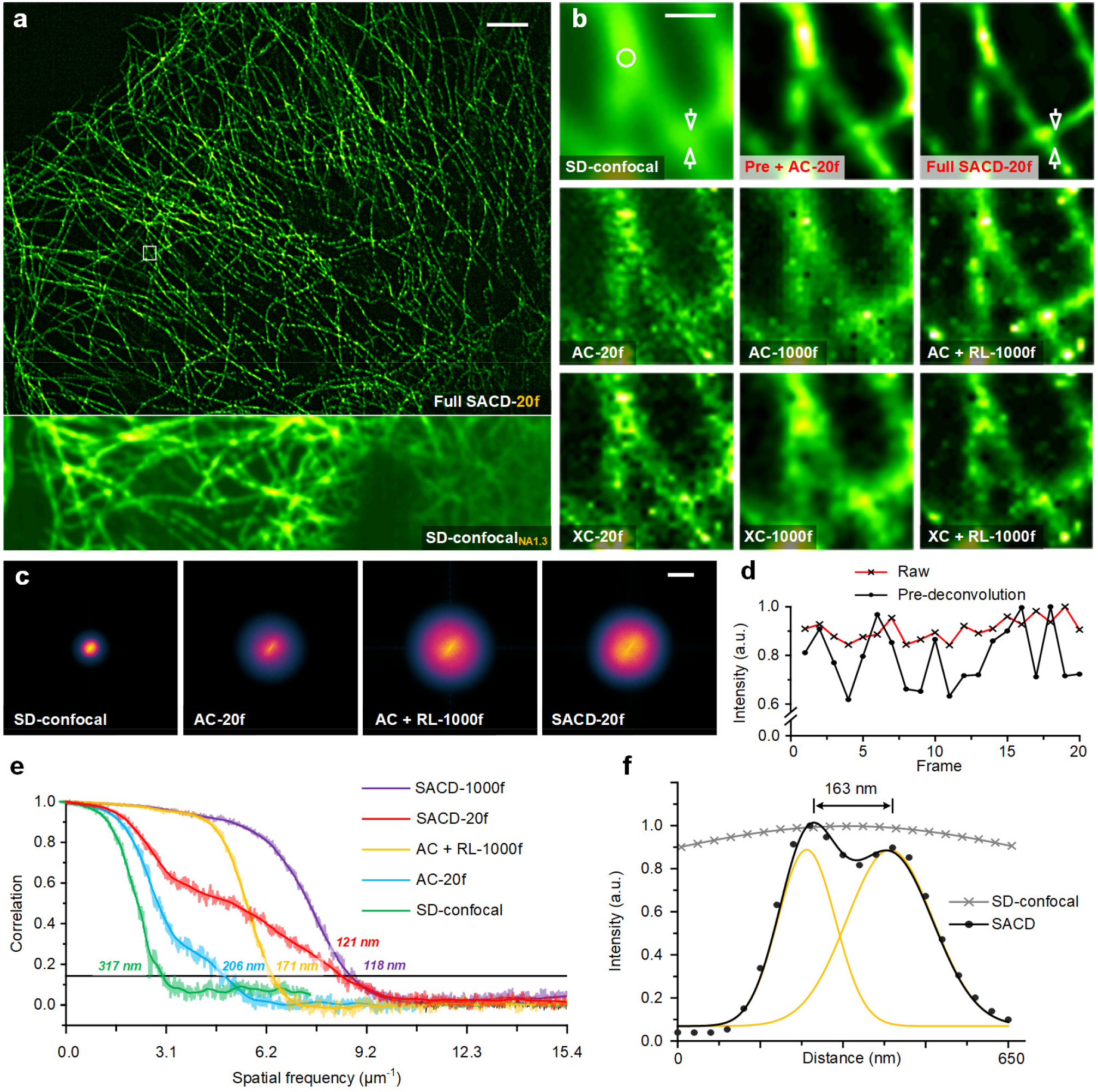
Evaluating SACD performance in 2D QD_525_ labeled microtubule versus SOFI. **(a)** A representative COS-7 cell labeled with QD_525_ under the SD-confocal and the SACD with 20 frames configurations was shown at the bottom and top sides. (**b**) Magnified views from the white box in (**a**) under different configurations. First row: SD-confocal, pre deconvolution followed by auto-correlation (AC) with 20 frames and full SACD reconstruction with 20 frames. Second row: pure AC with 20 frames, pure AC with 1000 frames, and AC with 1000 frames followed by RL deconvolution. Third row: pure cross-correlation (XC) with 20 frames, pure XC with 1000 frames, and XC with 1000 frames followed by RL deconvolution. (**c**) Fourier transforms of raw SD-confocal image, pure AC with 20 frames, AC with 1000 frames followed by RL deconvolution, and full SACD reconstruction with 20 frames. (**d**) Averaged intensity traces from pixels of the white circle in (**b**) for raw and raw after pre-deconvolution. (**e**) FRC analysis of the reconstructed images shows resolutions of 317 nm (raw SD-confocal), 206 nm (AC with 20 frames), 171 nm (AC with 1000 frames followed by RL deconvolution), 121 nm (SACD with 20 frames), and 118 nm (SACD with 1000 frames) respectively. (**f**) Intensity profiles and multiple Gaussian fitting of raw SD-confocal and 20-frame SACD for the structures of microtubule filaments indicated by the white arrows in (**b**). The numbers indicate the FWHM and distance between peaks. Details in **Supplementary Note 4**. Scale bars: (**a**) 5 μm; (**b**) 500 nm; (**c**) 10 μm.

**Extended Data Fig. 7.**
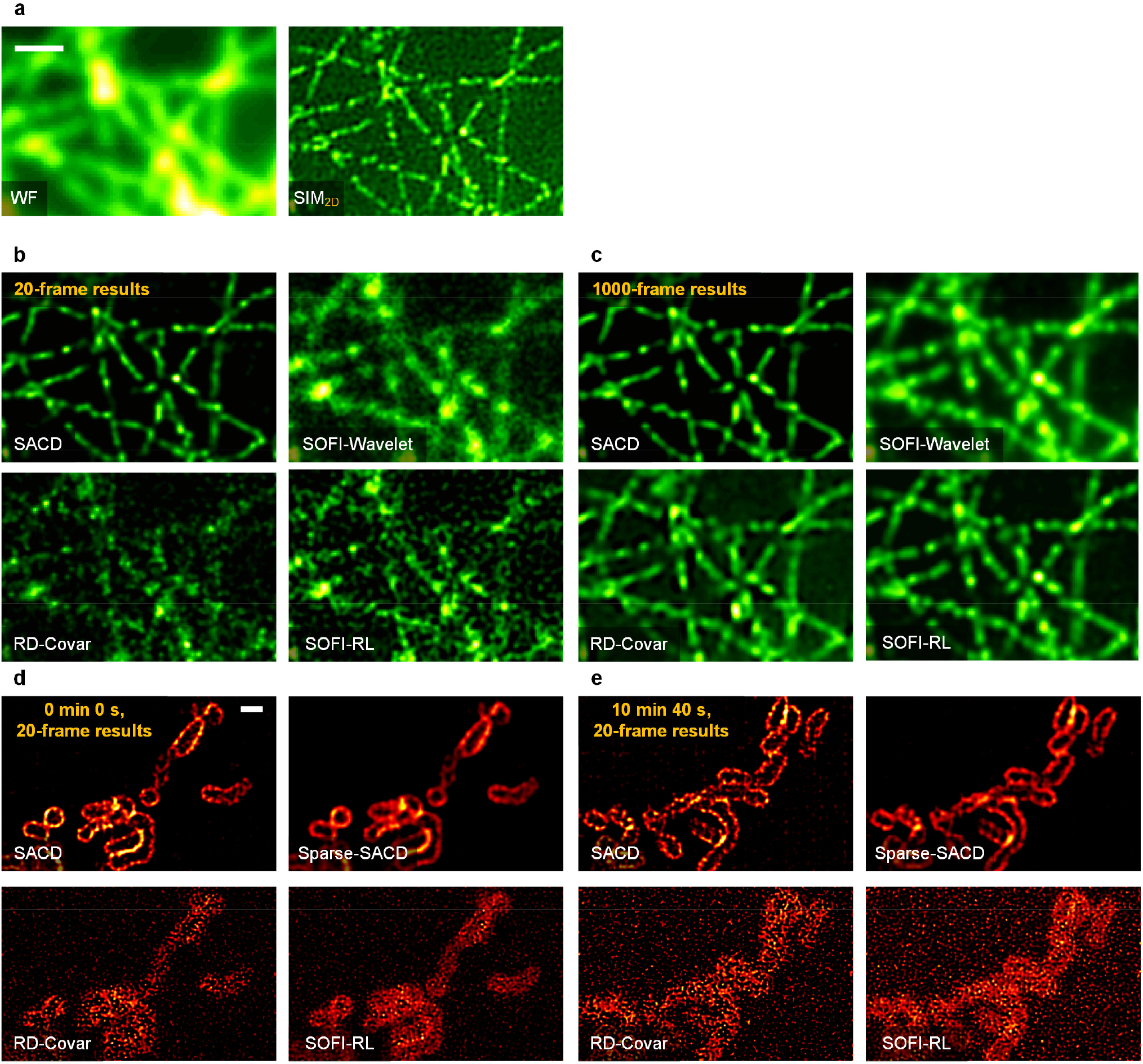
Experimental comparisons of different SOFI methods, SACD, and Sparse-SACD. (**a**) Microtubule filaments in a COS-7 cell labeled with QD525 imaged by the wide-field microscopy (left) and 2D-SIM (right) (*c*.*f*., **Fig. 1**). (**b, c**) Reconstruction results using 20 frames (**b**) and 1000 frames (**c**). (**b, c**) (**d, e**) Reconstruction results of live COS-7 cell labeled with Skylan-S-TOM20 imaged at 37°C imaged by SD-confocal (*c*.*f*., **Fig. 4d**) using 20 frames at time points 0 min 0 s (**d**) and 10 min 40 s (**e**). Details in **Supplementary Note 7**. Scale bars: 1 μm.

**Extended Data Fig. 8.**
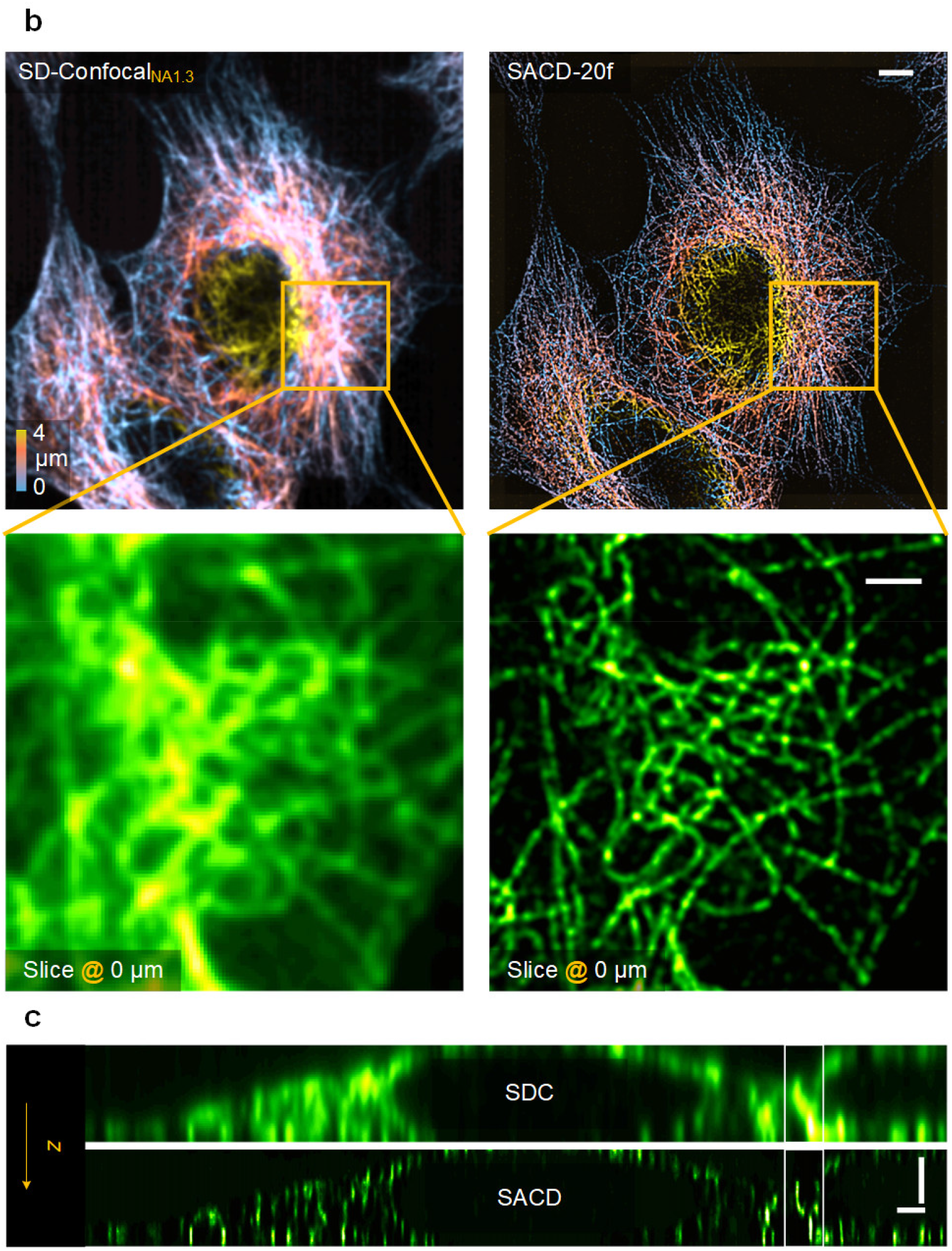
Magnified Fig. 2b and Fig. 2c for better visualization. (**b**) Color-coded three-dimensional distributions of microtubule filaments in a COS-7 cell labeled with QD_525_. Not overlapped view of SD-confocal (top left) and SACD (top right) images from **Fig. 2b**. The bottom panels show the *xy* slices (at 0 μm of *z* axial position) of SD-confocal (left) and SACD (right). (**c**) Corresponding vertical sections of SD-confocal (left) and SACD (right) from **Fig. 2c**. Scale bars: (**b**) 5 μm; (**b** inset, **c**) 2 μm.

**Extended Data Fig. 9.**
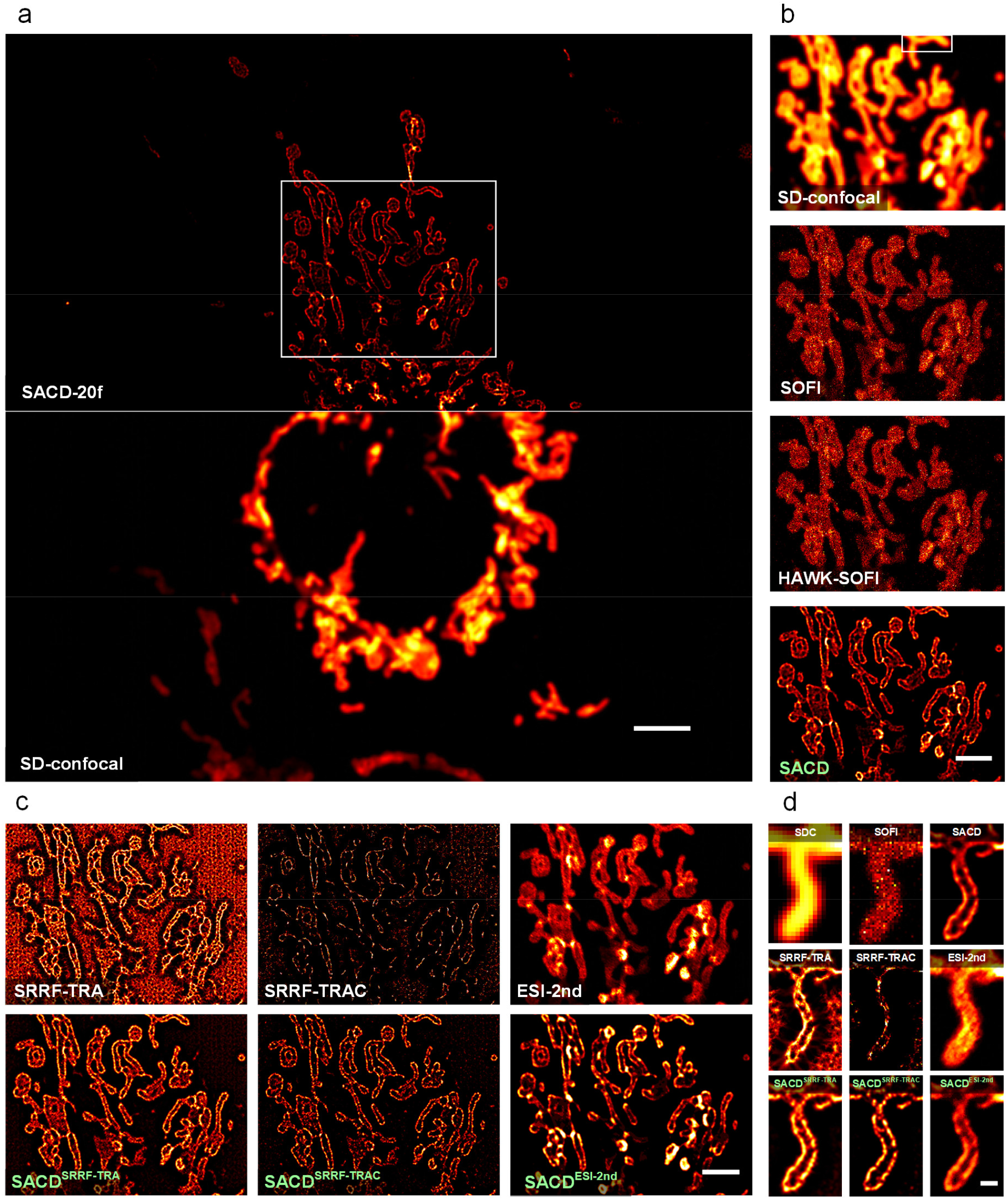
The flexibility of the SACD concept on other fluctuation-based SR techniques. (**a**) A representative live COS-7 cell labeled with Skylan-S-TOM20 imaged by SD-confocal (bottom) and SACD using 20 frames (top) (**Supplementary Video 3**). (**b**) Comparison of results (enlarged regions enclosed by the white box in **a**) of SD-confocal (1^st^ row, averaged by 20 frames), SOFI (2^nd^ row, reconstructed by 20 frames), HAWK-SOFI (3^rd^ row, reconstructed by 20 frames), and SACD (4^th^ row, reconstructed by 20 frames). (**c**) Comparison results (enlarged regions enclosed by the white box in **a**) of SRRF with temporal radiality average (SRRF-TRC, top left), SACD assisted SRRF-TRC (bottom left), SRRF with temporal radiality auto-correlation (SRRF-TRAC, top middle), SACD assisted SRRF-TRAC (bottom middle), ESI with second-order (ESI-2nd, top right), and SACD assisted ESI-2nd (bottom right). (**d**) Highlighted regions from the white box in (**b**). Details in **Supplementary Note 5**. Scale bars: (**a**) 5 μm; (**b**) 3 μm; (**c**) 500 nm.

**Extended Data Fig. 10.**
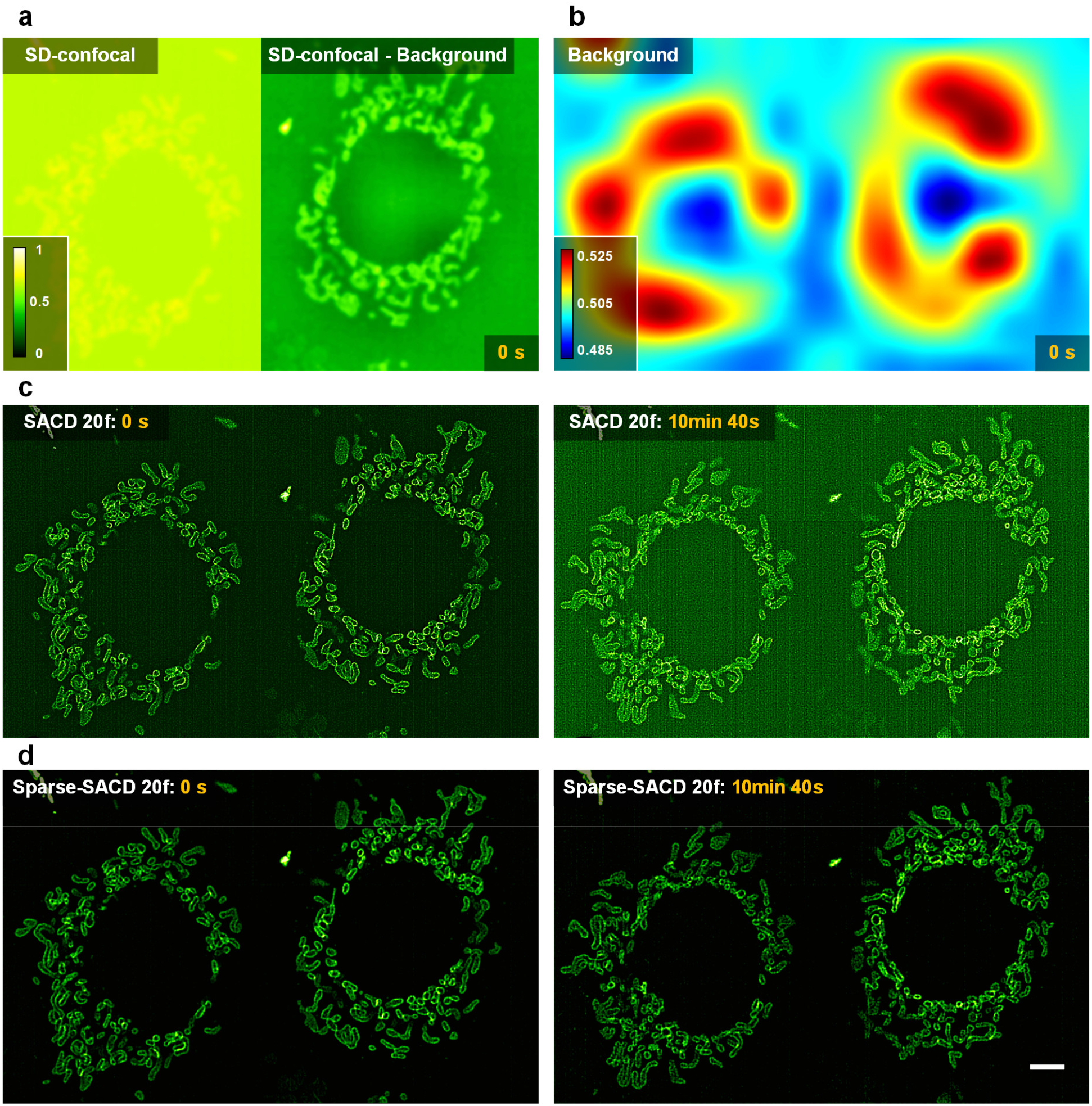
Sparse deconvolution assisted SACD for long-term live-cell SR imaging (*c*.*f*., Fig. 4c). (**a**) A representative live COS-7 cell labeled with Skylan-S-TOM20 imaged by SD-confocal (left) and SD-confocal after background subtraction (right). (**b**) The estimated background. (**c**) SACD results at time points 0 s (left) and 10 min 40 s (right). (**d**) Sparse-SACD results at time points 0 s (left) and 10 min 40 s (right). Scale bar: 5 μm.

