## Supplementary Information for "Enhancing detectable fluorescence fluctuation for high-throughput and four-dimensional live-cell super-resolution imaging"

### Content

**Supplementary Figures.** ... 3

**Supplementary Notes.** ... 8

Supplementary Note 1 | Characterizing properties of the pre-RL-deconvolution. ... 8

Supplementary Note 2 | Simulation comparisons of different SOFI methods and SACD. ... 10

Supplementary Note 3 | Benchmarking performances of different pre-processes. ... 14

Supplementary Note 4 | Experimental comparisons of SOFI and SACD. ... 15

Supplementary Note 5 | SACD-assisted diverse fluctuation-based SR microscopies. ... 17

Supplementary Note 6 | Simulation comparisons of SOFI, SACD, and Sparse-SACD under ultralow SNR. ... 19

Supplementary Note 7 | Experimental comparisons of different SOFI methods, SACD, and Sparse-SACD. ... 20

**Captions for Supplementary Movies.** ... 21

**References.** ... 27

### Supplementary Figures.

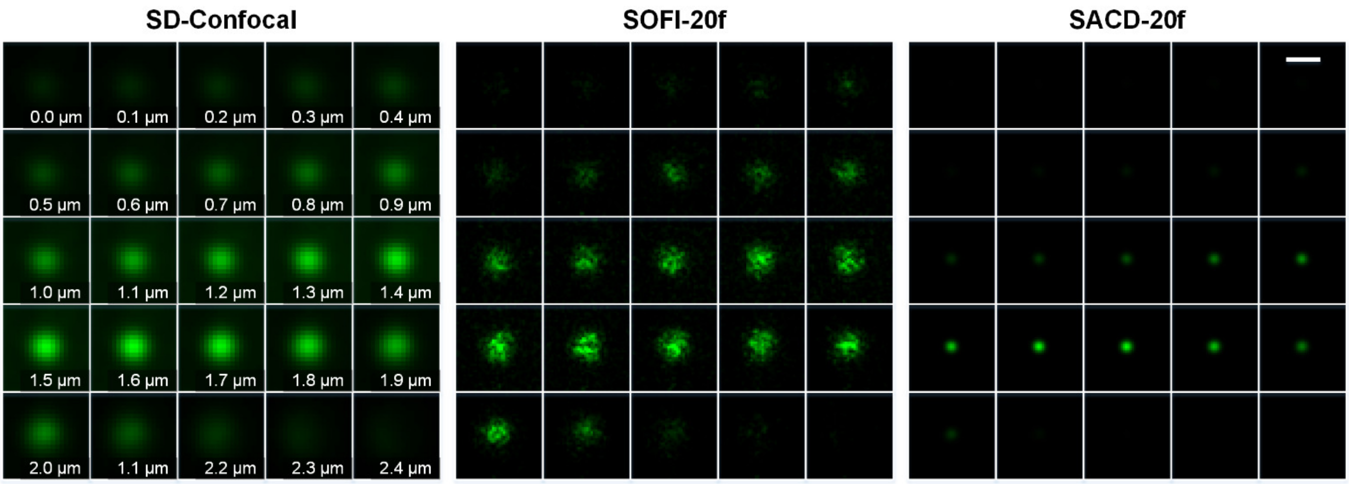


**Supplementary Fig. 1 | Full data representation of QD525 samples** (*c.f.*, **Fig. 2a**)**.** Axial montage of single $QD_{525}$ imaged by SD-confocal (left), SOFI using 20 frames (middle), and SACD using 20 frames (left). Scale bar: 500 nm.

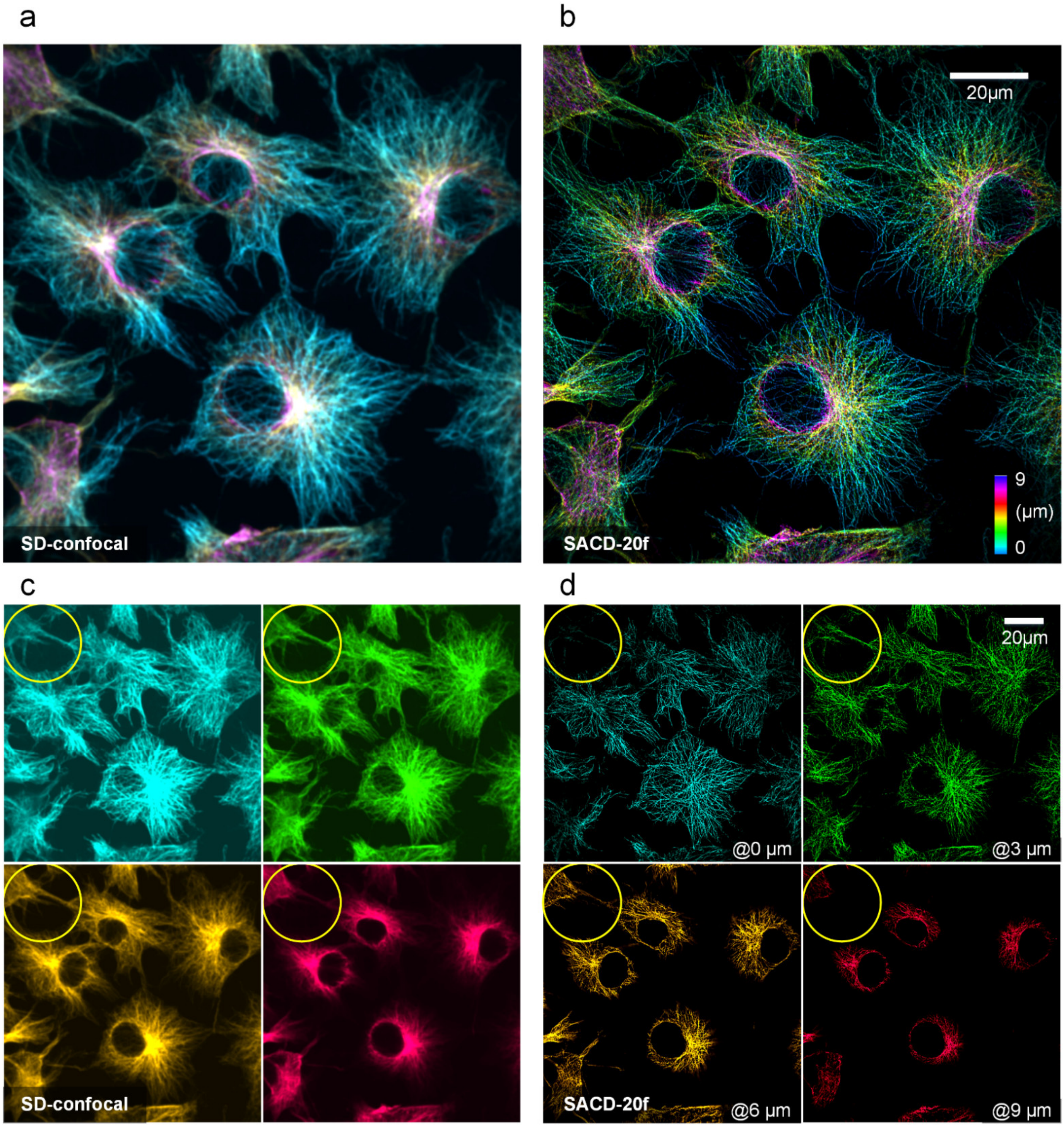


**Supplementary Fig. 2 | Full data representation of SACD large FOV 3D SR imaging.** (**a**, **b**) Large FOV, color-coded, three-dimensional distributions of microtubule filaments in COS-7 cells labeled with $QD_{525}$ (*c.f.*, **Fig. 2e**) imaged by the SD-confocal (**a**) and the SACD with 20 frames (**b**). (**c**, **d**) 4 color-coded axial slices under SD-confocal (**c**) and SACD (**d**) configurations at 0 µm, 3 µm, 6 µm, and 9 µm axial positions. The yellow circles indicate the axial resolution improvements by the SACD reconstruction. Scale bar: 20 µm.

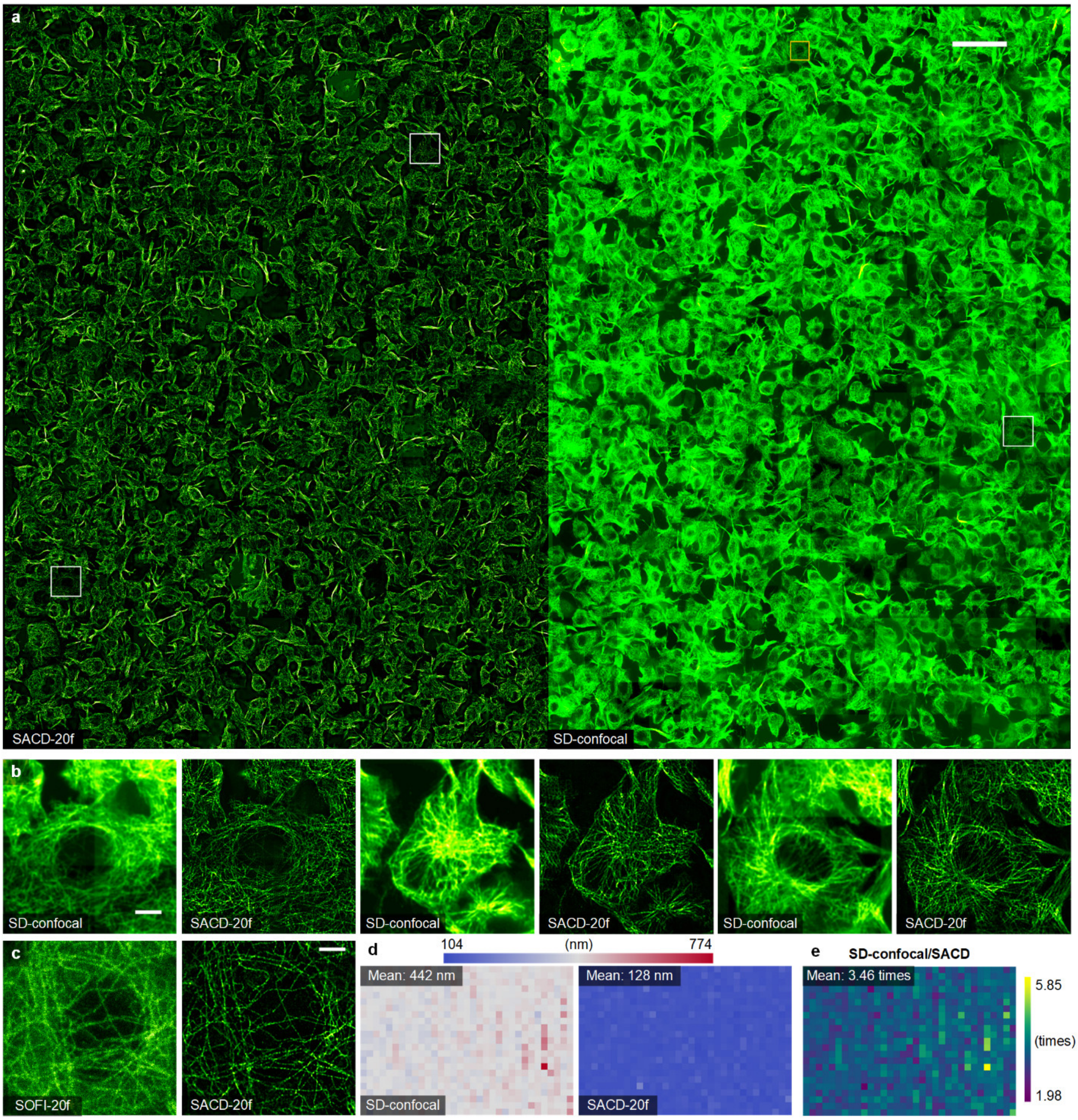


**Supplementary Fig. 3 | Full data representation of SACD high-throughput SR imaging (*c.f.*, Fig. 3). (a)** SACD reconstruction (left) and SD-confocal image (right) of a ~2.0 mm × 1.4 mm area. (**b**) Magnified views of the white-boxed regions in (**a**). (**c**) Enlarged views of the yellow boxed regions in (**a**). (**d**) The FRC resolution distribution over the entire FOV of SD-confocal (left) and SACD (right). (**e**) The result of the division of two FRC resolution maps from (**e**). Scale bars: (**a**) 100 μm; (**b**) 10 μm; (**c**) 5 μm.

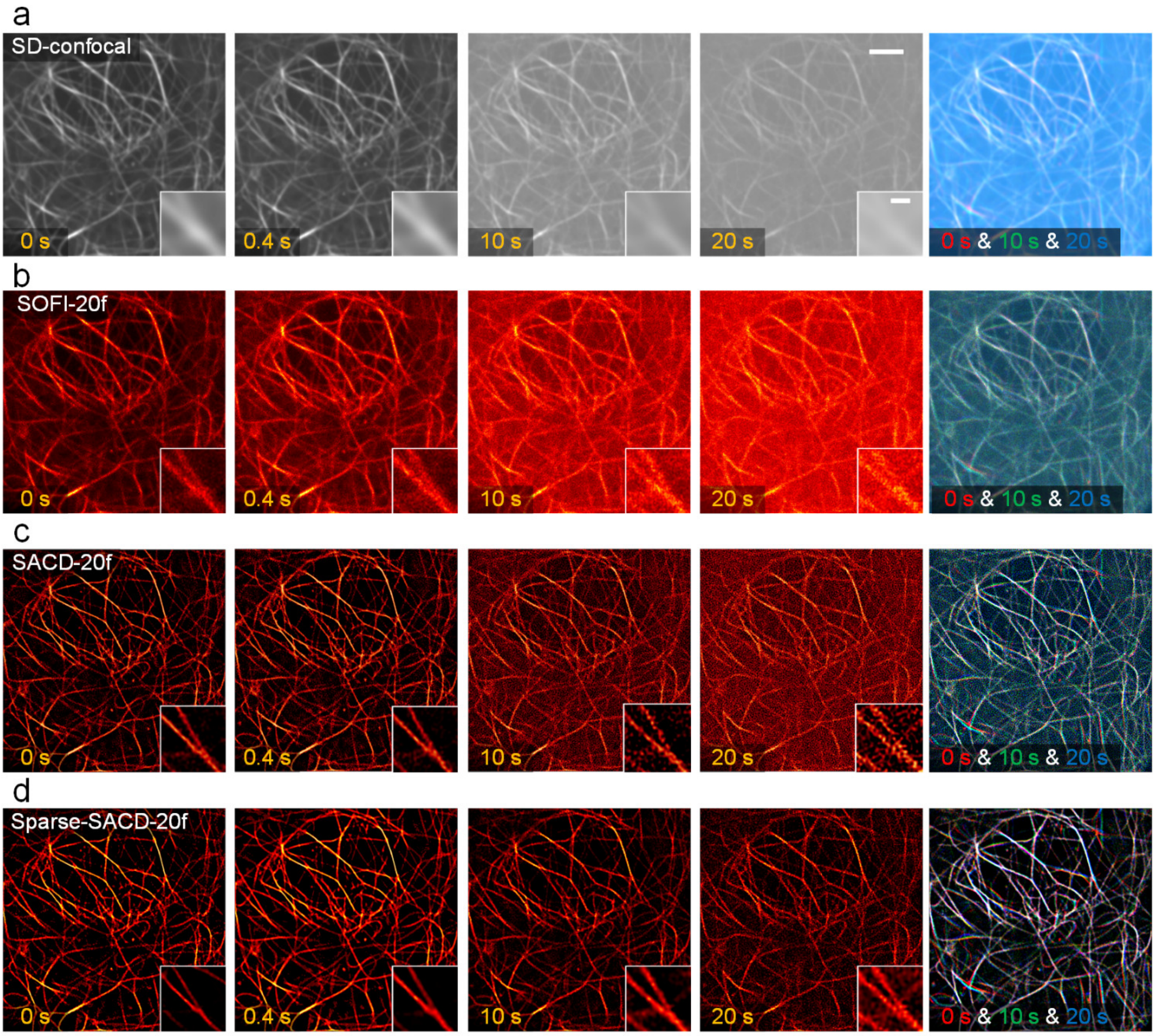


**Supplementary Fig. 4 | Sparse-SACD for low-SNR live-cell SR imaging (*c.f.*, Fig. 4b).** (**a-d**) The first four columns: A representative live COS-7 cell labeled with MAP4-Skylan-S imaged by SD-confocal (**a**), SOFI (**b**), SACD (**c**), and Sparse-SACD (**d**) at time points 0 s, 0.4 s, 10 s, and 20 s. The fifth column: Color-coded temporal projection of 0 s (red channel), 10 s (green channel), and 20 s (blue channel). Scale bars: 5 μm; inset 1 μm.

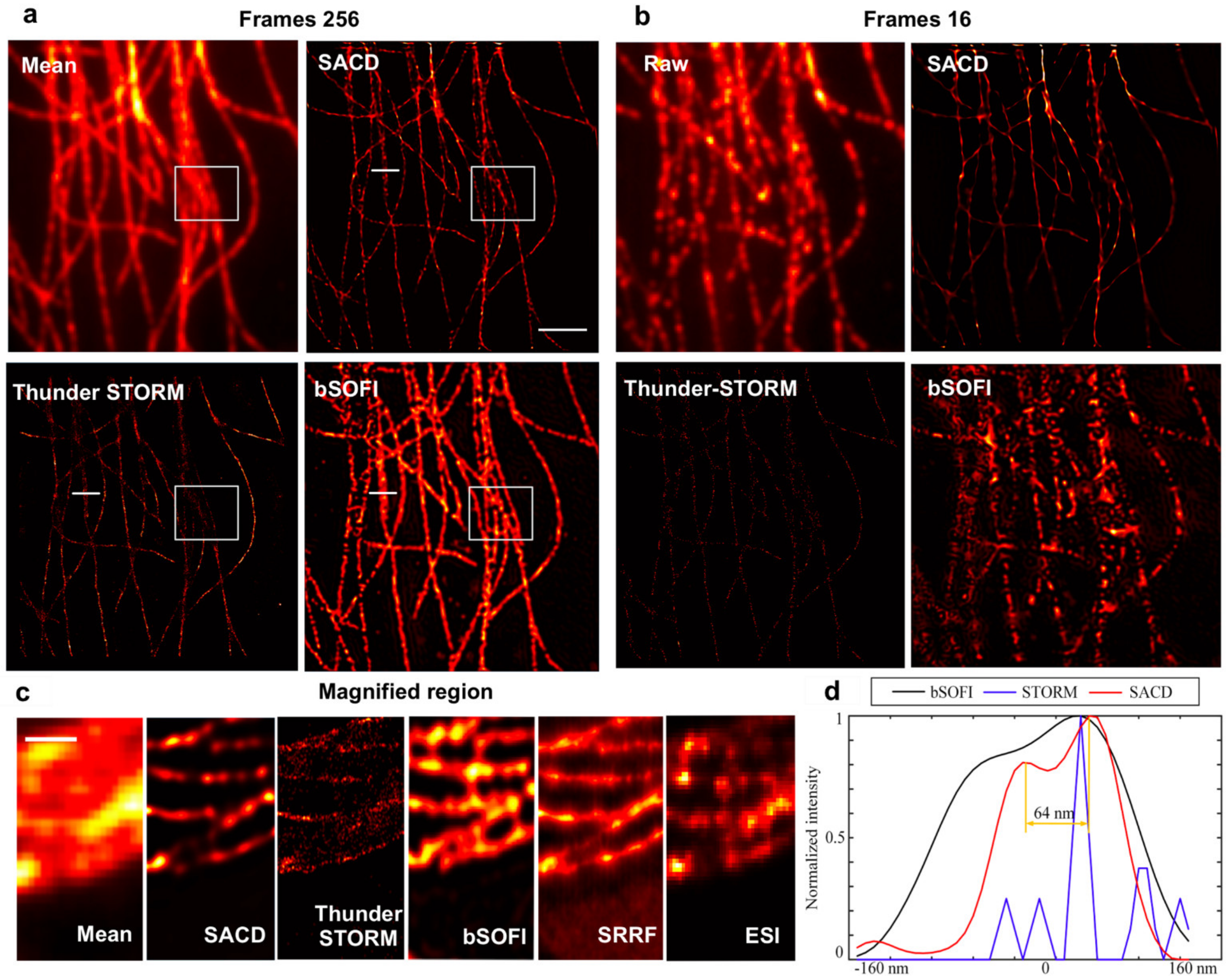


**Supplementary Fig. 5 | SACD result of the high-density SMLM image sequence and comparisons with the other SR methods.** (**a**, **b**) The TIRF image and the reconstructed images from the first 256 frames (dataset from ref[1]; 500 frames in total; NA as 1.3; frame rate as 25 f.p.s.; and the effective CCD pixel size is 100 nm). (**a**) The direct reconstructions from the first 256 frames. From top left to bottom right: Averaged TIRF image, SACD result (with 4$^{th}$ order), Thunder-STORM result, and 4$^{th}$ order bSOFI result. (**b**) The first 256 frames were averaged to create 16 higher-density frames (per 16 frames averaged as 1 frame) before reconstructions. From top left to bottom right: A single raw TIRF image, SACD result (with 4$^{th}$ order), Thunder-STORM result, and 4$^{th}$ order bSOFI result. (**c**) The magnified result of the white boxed region. (**d**) Profiles of the projection through the dotted lines in (**a**). Scale bars: (**a**) 2 µm; (**c**) 500 nm.

### Supplementary Notes.

#### Supplementary Note 1 | Characterizing properties of the pre-RL-deconvolution.

##### Supplementary Note 1.1 | RL deconvolution maintains the linear responses across frames.

The Richardson-Lucy (RL)[2, 3] deconvolution is regarded as a nonlinear deconvolution approach to iteratively recovering the underlying fluorescence signal, and its iterative formula in a product form commonly will cause an overall nonlinear intensity increase. To examine if this nonlinearity will influence the fluctuation signal along the temporal axis, we provided two representative cases by simulating the fluctuations of one molecule and two separative molecules (**Extended Data Fig. 2a-2d**). First, we created a single molecule with a cosine intensity fluctuation and convolved it with a 220 nm Gaussian PSF. In this case, we only included the Poisson noise (**Extended Data Fig. 2a**) on each frame and processed the resulting image stack with RL deconvolution. After acquiring the temporal responses of the pixel on the molecule center, we found the normalized temporal signals have perfectly identical distributions before and after RL deconvolution (**Extended Data Fig. 2b**), indicating that RL deconvolution can maintain the temporal linearity of the fluctuated signal. Second, we generated two blinking molecules with a distance slightly larger than the subsequently convolved PSF. In addition to Poisson noise, Gaussian noise and background baseline noise were added to the image (mixed noise) (**Extended Data Fig. 2c**). After RL deconvolution, the overall temporal distribution is highly aligned to the raw signal, and even has better on/off contrast ratio, referencing ground-truth response (left, **Extended Data Fig. 2d**). Next coming to the pixel between the two molecules, the reference tells that this pixel is the background with no fluorescence, and the intensity distribution after RL deconvolution perfectly matches this fact (right, **Extended Data Fig. 2d**). On the other hand, the raw signal still has a fluctuating response caused by the crosstalk of the nearby molecules. Overall, these two examples demonstrate that RL deconvolution can maintain the temporal response of fluctuating molecules when reducing their fluctuation crosstalk.

##### Supplementary Note 1.2 | RL deconvolution reduces pixel-level fluctuations and enhances molecular-level fluctuations.

Fundamentally, SOFI relies on the independent fluctuation behaviors of molecules rather than the pixel-level fluctuations induced by noise. To examine the effects of RL deconvolution on these two different fluctuations, we simulated two experiments (**Extended Data Fig. 2e-2h**). First, we created a single molecule and convolved it with a 220 nm Gaussian PSF before adding Poisson noise (**Extended Data Fig. 2e**). Compared to its noise-free higher resolution ground truth, as pointed out by black arrows, the Poisson noise-induced pixel-level fluctuations are successfully reduced after RL deconvolution (**Extended Data Fig. 2f**), and this may benefit the subsequent cumulant calculation, reducing the potential statistical uncertainty. Second, we simulated two molecules with opposite cosine intensity fluctuations convolved by a 220 nm Gaussian PSF before adding mixture noise (**Extended Data Fig. 2g**). To visualize their fluctuation behaviors, we color-coded these two molecules with magenta and green, respectively (**Extended Data Fig. 2g**). In addition to the involved mixed noise, the opposite distribution of fluctuations significantly compromised the effective on/off contrast ratio (**Extended Data Fig. 2h**). In stark contrast to the raw image sequence, intensity profiles after RL deconvolution are recovered back to the original fluctuation distributions (**Extended Data Fig. 2h**), and this may also facilitate the following calculation.

#### Supplementary Note 2 | Simulation comparisons of different SOFI methods and SACD.

There are several related SOFI methods developed to reduce the number of frames for reconstruction. In specific, the SOFI-wavelet[4] used a two-step wavelet-based spatiotemporal filter before SOFI calculation to enhance the SOFI efficiency. The SOFI-RL[5, 6] involved a post-deconvolution to further enhance the resolution and contrast. The RD-Covar[7] introduced a covariance-based SOFI calculation to enhance the SOFI efficiency and post-deconvolution to further enhance the resolution and contrast. However, these methods are sensitive to either background noise and SNR, or multiple tunable parameters, and cannot achieve acceptable performance under the 20-frame configuration. In this section, we quantitatively compared the performances of SACD and these SOFI methods under different conditions using two typical simulated structures. We summarized that only SACD can achieve superior performance in both fidelity and uncertainty under the 20-frame configuration and is insensitive to SNR conditions.

**Ring structures.** First, we created ring structures of different diameters (100 nm to 420 nm, with a 40 nm interval) on a 10 nm grid. Then, following ref.[8], we generated temporal blinking sequences of the fluorescence emitters and convolved them with a 220 nm PSF subsequently (**Methods**). The resulting image stack was downsampled 6 times (pixel size 60 nm). To synthesize more realistic experimental imaging conditions, the cytosol out-of-focus background and mixture noise under three different levels ('*low*', '*medium*', and '*high*') were included in the images (**Extended Data Fig. 5b**). We used the three modified SOFI methods mentioned above and SACD to reconstruct images from the 20-frame raw stack. First, under a visual examination of different methods in **Extended Data Fig. 5c-5e**, the image fidelity of SOFI-wavelet, SOFI-RL, and RD-Covar dramatically decreases with the deterioration of imaging conditions, while the SACD is robust (**Extended Data Fig. 5f**), consistently reconstructing structures without amplifications of artifacts.

Because one two-peak analysis may be not sufficient to validate that the ring is fully resolved[9], to quantitatively evaluate the contrast across the entire ring structure, we proposed a ring quality ratio (RQR) criterion (**Extended Data Fig. 5h**). To avoid overconfident determination, we chose a value 0.4, much larger than the Raleigh criterion (0.1801), as the threshold to determine whether the rings are resolved successfully. According to this criterion, under the *high* SNR condition, SOFI-wavelet, SOFI-RL, and RD-Covar reached a limitation at resolving 300 nm, 260 nm, and 180 nm, respectively, while the SACD could separate rings at 140 nm apart. Under the *medium* and *low* SNR conditions, the SACD could reconstruct the 180 nm ring, referring to the ~110 nm resolution ground-truth image (**Extended Data Fig. 5a**), while other SOFI methods distinguished a ring with a minimum diameter of 220 nm. Compared to the other two SOFI methods, the RD-

Covar has the potential to achieve high-resolution results with the 20-frame configuration but is extremely sensitive to noise conditions. Even under our high SNR condition, the outlines of RD-Covar reconstructed rings are insufficiently continuous. On the other hand, our SACD achieved high-fidelity resolution enhancements under various conditions.

To evaluate the fidelity across different numbers of frames adopted, we calculated the structural similarity (SSIM)[10] (**Extended Data Fig. 5g**) values against the ground truth of different methods, and we also included the standard deviation (STD)[11] (**Extended Data Fig. 5i**) values reflecting the reconstruction uncertainty. We found a rapid increase in the fidelity and stability of SACD reconstructions from 5 to 20 frames used, and after 20 frames, the SSIM and STD values reached a plateau. In contrast, SSIM and STD values of the other methods continued to vary as the used number of frames rose and did not stop even when the number of frames reached 1000. These results demonstrate that the 20-frame reconstruction is adequate for SACD in both fidelity and stability.

**Filaments.** We repeated the verification using microtubule-like filament structures (**Supplementary Fig. 6**). As expected, we found a similar distribution of fidelity on these reconstructions of filament structures (**Supplementary Fig. 6g**). During the decrease of SNR condition, the SOFI-wavelet, SOFI-RL, and RD-Covar suffered from the dominating background over the entire object area, and amplified strong snow-flake artifacts (**Supplementary Fig. 6c-6e**). In contrast, the SACD results are impressed with sturdily recognizing and reconstructing the filament structures without fidelity degradation (**Supplementary Fig. 6f**).

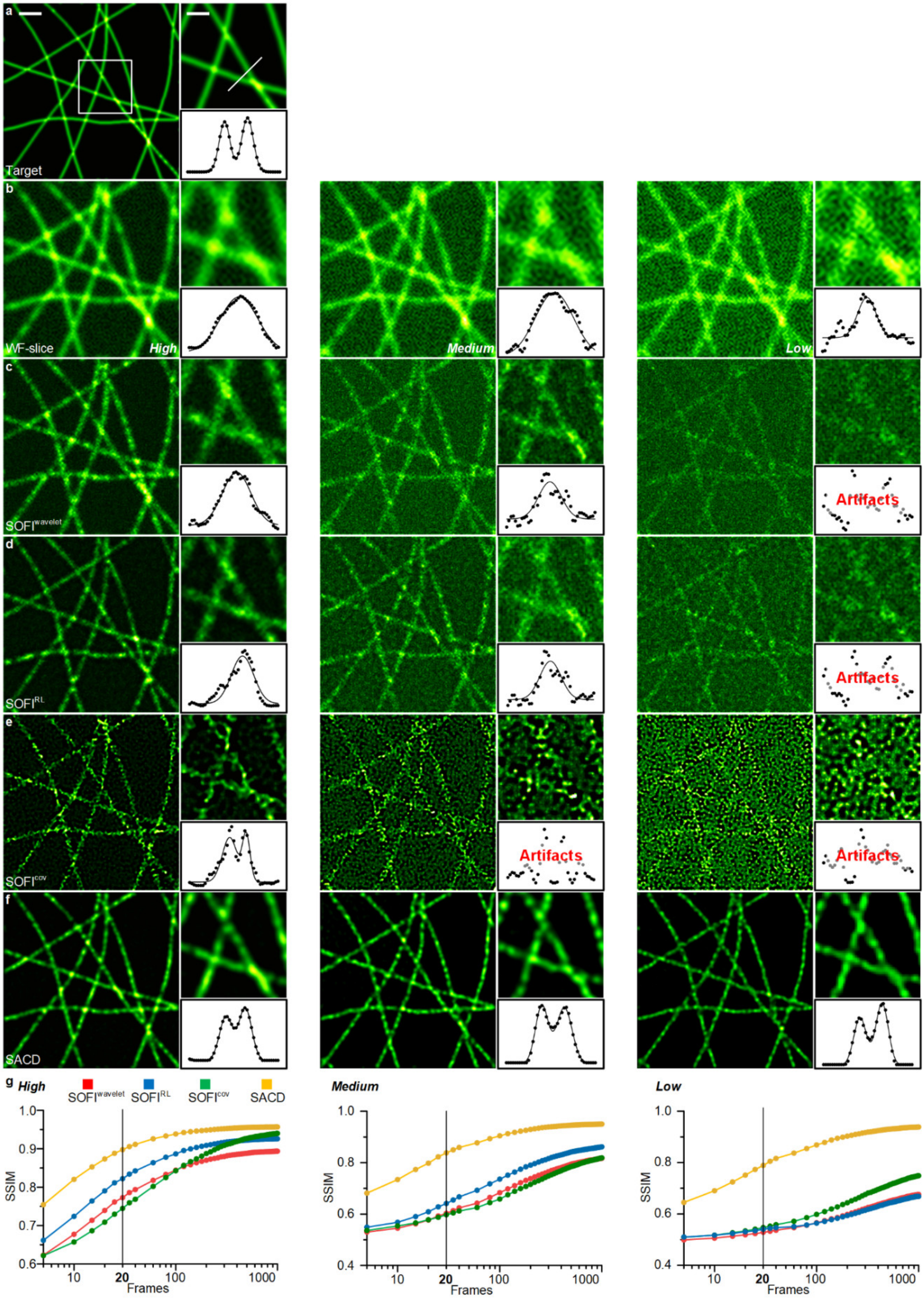

a
Target
b
WF-slice
High
Medium
Low
c
SOFIwavelet
Artifacts
d
SOFIRL
Artifacts
e
SOFIcov
Artifacts
Artifacts
f
SACD
g
High
SOFIwavelet
SOFIRL
SOFIcov
SACD
Medium
Low
SSIM
Frames


143

**Supplementary Fig. 6 | Simulation comparisons of different SOFI methods and SACD under different SNRs using filament structures.** (**a**) The synthetic filament structures were convolved with a 110 nm PSF (pixel size 20 nm) as the ground truth. (**b-f**) Wide-field (WF, **b**) raw images and SOFI-wavelet (**c**), SOFI-RL (**d**), RD-Covar (**e**), and SACD (**f**) reconstructions using 20-frame under '*High*' (right), '*Medium*' (middle), and '*Low*' (left) SNR conditions. The structures were convolved with a 220 nm PSF, and down-sampled 3 times before adding the cytosol background, out-of-focus light, Poisson noise, Gaussian readout noise, and baseline background under three different levels to be the raw WF image stacks. The sub-images at the upper right corner are the enlarged view from the white box. The sub-images at the lower right corner are the intensity profiles and multiple Gaussian fitting from the white line. (**g**) The SSIM curves of different methods under '*High*' (right), '*Medium*' (middle), and '*Low*' (left) SNR conditions. Scale bars: (**a**) 500 nm; (**a**, inset) 100 nm.

#### Supplementary Note 3 | Benchmarking performances of different pre-processes.

To examine the performance differences of various pre-processes, we replaced our pre-RL-deconvolution with a low pass filter or a Wiener deconvolution and used the same dataset from **Extended Data Fig. 5** and **Supplementary Fig. 6** under the *low* SNR condition. It can be seen in **Supplementary Fig. 7**, the low pass pre-filter[12] can eliminate strong noise components and enhance the continuity of SOFI-RL on both ring (SSIM as 0.62) (**Supplementary Fig. 7c**) and filament (SSIM as 0.65) structures (**Supplementary Fig. 7g**). However, this pre-filter will also compromise the achievable resolution, leading to relatively blurry results compared to SACD (SSIM as **0.88** or **0.81**) (**Supplementary Fig. 7b**, **7f**). On the other hand, the reconstruction of Wiener deconvolution is sensitive to the damping coefficient, and its optimal quality (SSIM as 0.67 or 0.66) is still not comparable to SACD (SSIM as 0.88 or 0.81) (**Supplementary Fig. 7d**, **7h**). As a result, we can conclude that our SACD reconstruction process is the optimal solution for fluctuation-based SR microscopy.

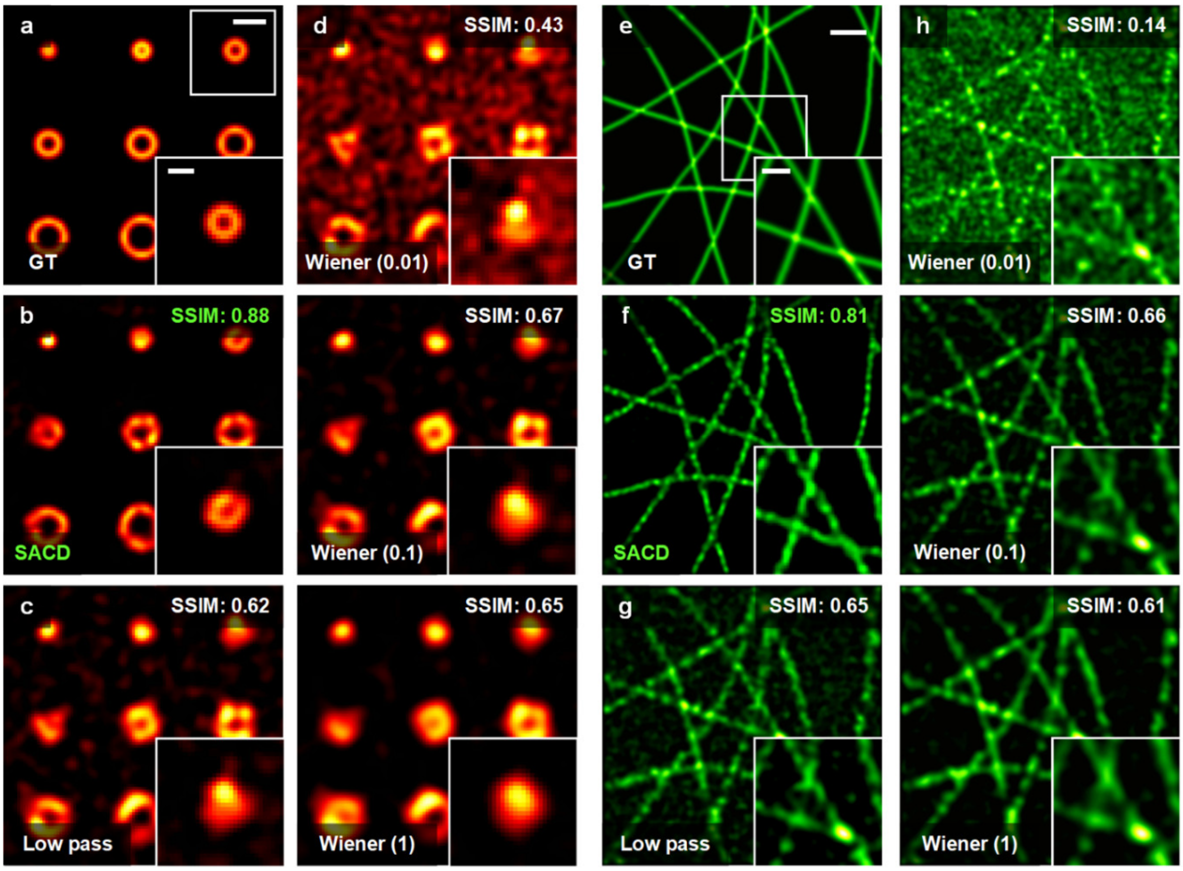


**Supplementary Fig. 7 | Comparisons of different pre-processes under the *low* SNR condition.** (**a-h**) Comparisons using ring (**a-d**) (*c.f.*, **Extended Data Fig. 5**) and filament (**e-h**) (*c.f.*, **Supplementary Fig. 6**) structures. (**a**, **e**) Ground truth images. (**b**, **f**) SACD results. (**c**, **g**) Reconstructions with replacing pre-RL-deconvolution to a low pass filter (using the cutoff frequency of optical transfer function). (**d**, **h**) Reconstructions with replacing pre-RL-deconvolution to a Wiener deconvolution using 0.01 (top), 0.1 (middle), and 1 (bottom) as Wiener damping coefficients. The sub-images at the lower right corner are the enlarged view from the white box in (**a**) and (**e**). Scale bars: (**a**) 300 nm; (**a**, inset) 150 nm; (**e**) 500 nm; (**e**, inset) 100 nm.

**Supplementary Note 4 | Experimental comparisons of SOFI and SACD.**

Using a commercial spinning disk confocal (SD-confocal) microscope equipped with an objective (numerical aperture, NA = 1.3), we directly image the quantum dot 525 ($QD_{525}$) labeled fixed COS-7 cells (**Methods**, **Extended Data Fig. 6a**), in which 2000 frames were consecutively recorded. First, in **Extended Data Fig. 6b**, we compared the auto-correlation cumulant (AC)[13, 14] results using 20 frames. Apparently, by involving the pre-deconvolution, the 'Pre + AC-20f' result outperformed the one in 'AC-20f'. It is notable that even employing 1000 frames, the pure auto-correlation cumulant result represented an inferior quality. Likewise, combing the post-deconvolution, the image quality of the full SACD result ('Full SACD-20f') appeared even better than the result in full SOFI, in which the AC was followed by an RL deconvolution (AC + RL-1000f). When tracing the averaged fluorescence intensity inside the white circle in **Extended Data Fig. 6b**, as demonstrated in **Extended Data Fig. 6d**, it is found that the pre-deconvolution process significantly improved the detectable fluorescence fluctuation behavior (the effective on/off ratio) existing in such datasets. It indicates that the pre-deconvolution is the key point offering the following auto-correlation cumulant reconstruction high-performance only by fewer frames.

Also, in Fourier space (**Extended Data Fig. 6c**), it is observed that the 'SACD-20f' notably extended the effective optical transfer function (OTF), which appeared even better than the 'AC + RL-1000f' result. Better separation of intertwining microtubule filaments (**Extended Data Fig. 6f**) can be visualized in SACD due to the increased spatial resolution, which is also verified by FRC resolution analysis (**Extended Data Fig. 6e**). The original FRC resolution of SD-confocal is estimated as ~317 nm, and our SACD effectively doubled it reaching ~121 nm. Predictably, the 'AC-20f' led to a limited resolution improvement to ~206 nm due to the insufficient contrast ratio in real experiments. Even recording a large raw image sequence for 1000 frames, SOFI is hard to fully attain the expected doubled resolution, as seen in **Extended Data Fig. 6e**, in which it reached ~171 nm. Remarkably, the resolutions represented an unapparent difference between the 20 frames and 1000 frames in SACD images, which are ~118 nm and ~121 nm.

Since the signal was captured by a sCMOS camera (**Methods**), the shot noise affected the continuity and homogeneity of pure AC reconstruction to a large extent[5, 12]. As can be seen in **Extended Data Fig. 6b** (AC-20f), such shot noise is prone to induce artifacts and discontinuity on the fine structures of microtubule filaments. Therefore, we also involved the spatiotemporal cross-cumulant (XC)[5] as a further comparison. Considering the shot noise from different pixels being not correlated, we can benefit from the form of cross-cumulant, enabling the elimination of such shot noise[5, 12]. On the other hand, this approach highly depends on the exact knowledge of the PSF and the specific cross-correlation geometry for calculation, which is hard to optimize and more complex in computation compared to the AC method. Nevertheless, to examine the full ability of SACD, we still provided the results of XC calculated by 20 frames and 1000 frames, and the 'XC-1000f' result is also followed with a post-deconvolution. Among all these results in **Extended Data Fig. 6b**, we can see that the 'SACD-20f' not only reduces the necessary number of the frames adopted, but also eliminates the shot-noise for the AC method incidentally, achieving a superior image quality and resolution enhancement even against the XC calculation.

**Supplementary Note 5 | SACD-assisted diverse fluctuation-based SR microscopies.**

Heuristically, our SACD concept utilizes a pre-RL-deconvolution operation to optimize the detectable fluctuation behavior, and this feature may also benefit other fluctuation-based techniques for increasing their temporal resolvability as well. Initially, we used an open-source live-cell dataset (GFP-tagged microtubules in live HeLa cells imaged by TIRF mode[15], **Methods**, **Supplementary Fig. 8**) to evaluate the potential enhancement of the SACD cooperating with the SRRF algorithm[15]. The microtubule structures are overwhelmed by the excessive background in the 20-frame SRRF image to a large extent (**Supplementary Fig. 8e**). In contrast, the SACD-assisted SRRF, i.e., pre-RL-deconvolution followed by the SRRF reconstruction, is capable of revealing the fully representative microtubule filaments ($\mathrm{SACD}^{\mathrm{SRRF}}$, **Supplementary Fig. 8f**). By increasing the number of frames adopted, the 200-frame SRRF attained a comparable quality of reconstruction (**Supplementary Fig. 8d**). However, it is notable that such a long frame sequence configuration will induce considerable motion blur in the results (white arrows in **Supplementary Fig. 8j**), significantly reducing the efficient spatial resolvability.

To fully test the broad applicability of the SACD concept in assisting other fluctuation-based approaches, we examined a series of fluctuation-based results, including SOFI[13], SRRF, and ESI[16] in live COS-7 cells labeled by Skylan-S-TOM20. As shown in **Extended Data Fig. 9b**, the 20-frame SOFI failed to resolve the fine structures of outer mitochondrial membranes (OMM), while the Haar wavelet kernel pre-operation[17] (HAWK-SOFI) can scarcely provide any improvements on its resolvability. As expected, SACD revealed the tori cross-section of OMM genuinely (**Extended Data Fig. 9a**, **Extended Data Fig. 9b**, **Supplementary Video 3**). Another two frequently-used modalities have been employed and analyzed in **Extended Data Fig. 9c**, which are SRRF under two configurations, i.e., temporal radiality average (SRRF-TRC) and temporal radiality auto-correlation (SRRF-TRAC), and ESI as well. As highlighted in **Extended Data Fig. 9c**, we confirmed that our designed pre-RL-deconvolution is not only suitable for SOFI, but also SRRF and ESI. Focusing on a small region, a single mitochondrion in **Extended Data Fig. 9d**, SACD-assisted methods are impressed with sturdily recognizing and reconstructing the OMM structures, where the original SRRF is inclined to produce over-slimming 'line-like' OMM structures due to using a limited dataset. On the other side, both SOFI and ESI show their inadequate resolving capability and even suffer from the dominating artifacts over the entire object area. Likewise, these issues are thoroughly settled assisted by SACD, enabling background-free and high-fidelity SR imaging.

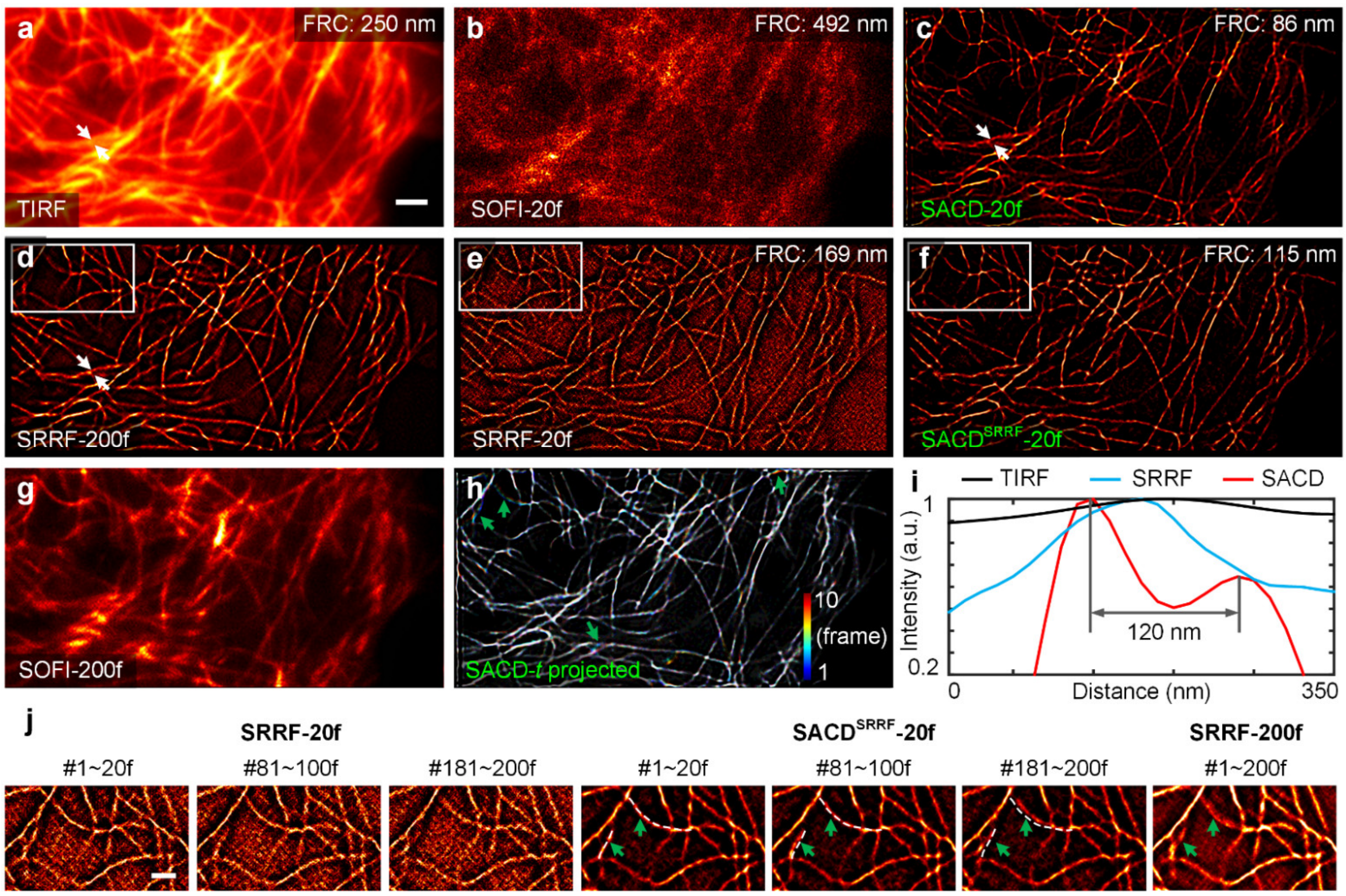


**Supplementary Fig. 8 | SACD and SACD-assisted SRRF reconstructions for 2D live-cell imaging.** (**a-g**) GFP-tagged microtubules in a live HeLa cell. Dataset from ref[15]. Raw TIRF image (**a**), 20-frame SOFI (**b**), 20-frame SACD (**c**), 200-frame SRRF (**d**), 20-frame SRRF (**e**) 20-frame SACD-assisted SRRF (SRRF with pre-deconvolution) (**f**) and 200-frame SOFI (**g**). (**h**) Color-coded temporal projections of microtubules within 10 SR frames (200 raw frames in total). The obvious motions are pointed by green arrows. (**i**) Intensity profiles indicated by white arrows in (**a**), (**c**), and (**d**) for TIRF (black), 200-frame SRRF (blue), and SACD (red), respectively. (**j**) Magnified views from white boxes in (**d-f**). The views reconstructed by 1~20 frames, 81~100 frames, and 181~200 frames for 20-frame SRRF and 20-frame SACD-assisted SRRF are shown on the left and the middle. The view reconstructed by 1~200 frames for 200-frame SRRF is represented on the right. The motion blur contained in 200-frame SRRF is pointed by the green arrows. As indicated by white dashed lines and green arrows in 20-frame SACD-assisted SRRF, such motion blur can be reduced effectively by the 20-frame reconstruction. Scale bars: (**a**) 2 μm; (**j**) 1 μm.

#### Supplementary Note 6 | Simulation comparisons of SOFI, SACD, and Sparse-SACD under ultralow SNR.

Although insensitive to noise levels, the continuity and fidelity of SACD will reduce when the SNR is lower to a certain extent. Thus, we next intend to test the differences between SOFI-RL (SACD without pre-RL deconvolution), SACD, and Sparse-SACD under more harsh imaging conditions. As can be seen in **Supplementary Fig. 9**, under such SNR, the SOFI-RL failed to reconstruct the ring (SSIM as 0.08) or filament structures (SSIM as 0.06). The failures of SOFI-RL were partially compensated by the pre-RL-deconvolution, and SACD (SSIM as 0.83 or 0.62) successfully resolved fine structures but the severe noise of raw data still induced background artifacts and continuity reduction. By replacing post-RL-deconvolution with Sparse deconvolution[9] constrained by sparsity and continuity prior knowledge, the Sparse-SACD (SSIM as **0.94** or **0.83**) results exhibit clear background and structures with high-fidelity.

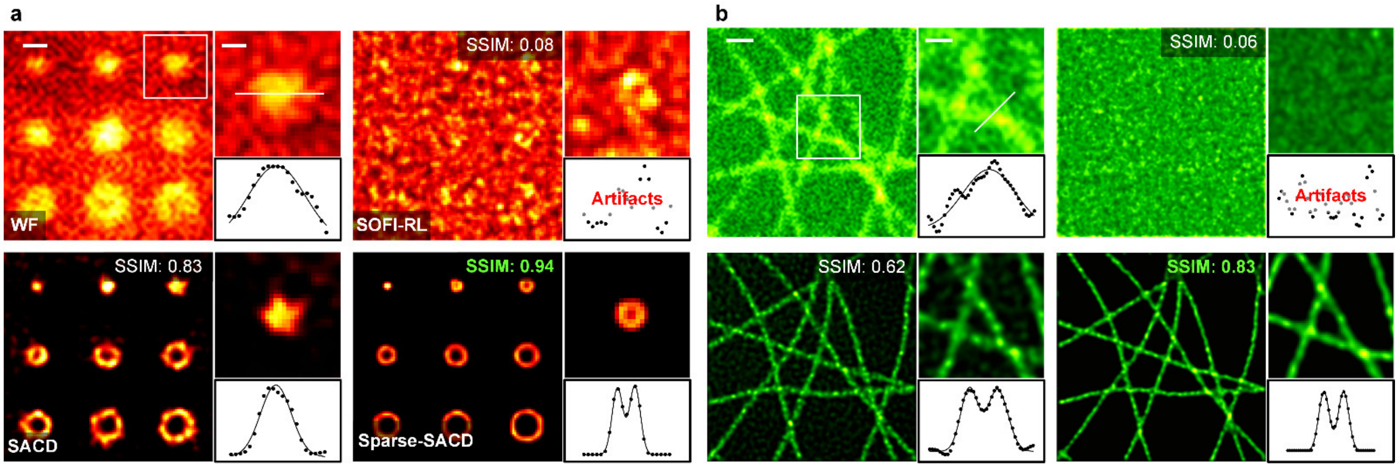


**Supplementary Fig. 9 | Comparisons of SOFI, SACD, and Sparse-SACD under *ultralow* SNR condition.** (**a**, **b**) Comparisons using ring (**a**) and filament (**b**) structures. First row: wide-field raw image (WF, left), and SOFI-RL result (right); Second row: SACD (left) and Sparse-SACD (right) results. The sub-images at the upper right corner are the enlarged view from the white box. The sub-images at the lower right corner are the intensity profiles and multiple Gaussian fitting from the white line. Scale bars: (**a**) 300 nm; (**a**, inset) 150 nm; (**b**) 500 nm; (**b**, inset) 100 nm.

#### Supplementary Note 7 | Experimental comparisons of different SOFI methods, SACD, and Sparse-SACD.

We next intend to examine the performance of SACD against other modified SOFI methods under a wide-field microscope using fixed cells with 2D-SIM reference[18] (**Extended Data Fig. 7a**). Under 20-frame configuration, the SOFI-wavelet, SOFI-RL, and RD-Covar cannot decode the fine structures of microtubules from molecule fluctuations, and in contrast SACD successfully reconstruct the cytoskeleton network, perfectly matching to the 2D-SIM ground truth (**Extended Data Fig. 7b**). When using 1000 frames, these three modified SOFI methods achieved sufficient performance for resolving intertwining filaments, and on the other hand SACD exhibited an almost identical result comparing to the 20-frame result (**Extended Data Fig. 7c**). This indicates that our SACD massively improve the SOFI sufficiency, and the 20-frame configuration is adequate for high-quality SR imaging.

Recording sensitive processes of outer mitochondrial membranes (OMM) using a protein label with low excitation intensity will induce strong readout noise, which translated into the reconstruction of severely aberrated SR images using the modified SOFI methods (SOFI-RL and RD-Covar), especially after long-term imaging (~10 min) (**Extended Data Fig. 7d**, **7e**). In contrast, our SACD consistently recovered the hollow structures of OMM. However, as the average fluorescence (at 10 min 40 s) progressively decreased to ~50% of its initial value (at 0 min 0 s) during the recording, artifacts in SACD images also increased (**Extended Data Fig. 7e**). The addition of sparsity and continuity constraints (Sparse-SACD) suppressed the background artifacts and largely improved the structural continuity (**Extended Data Fig. 7e**).

#### Captions for Supplementary Movies.

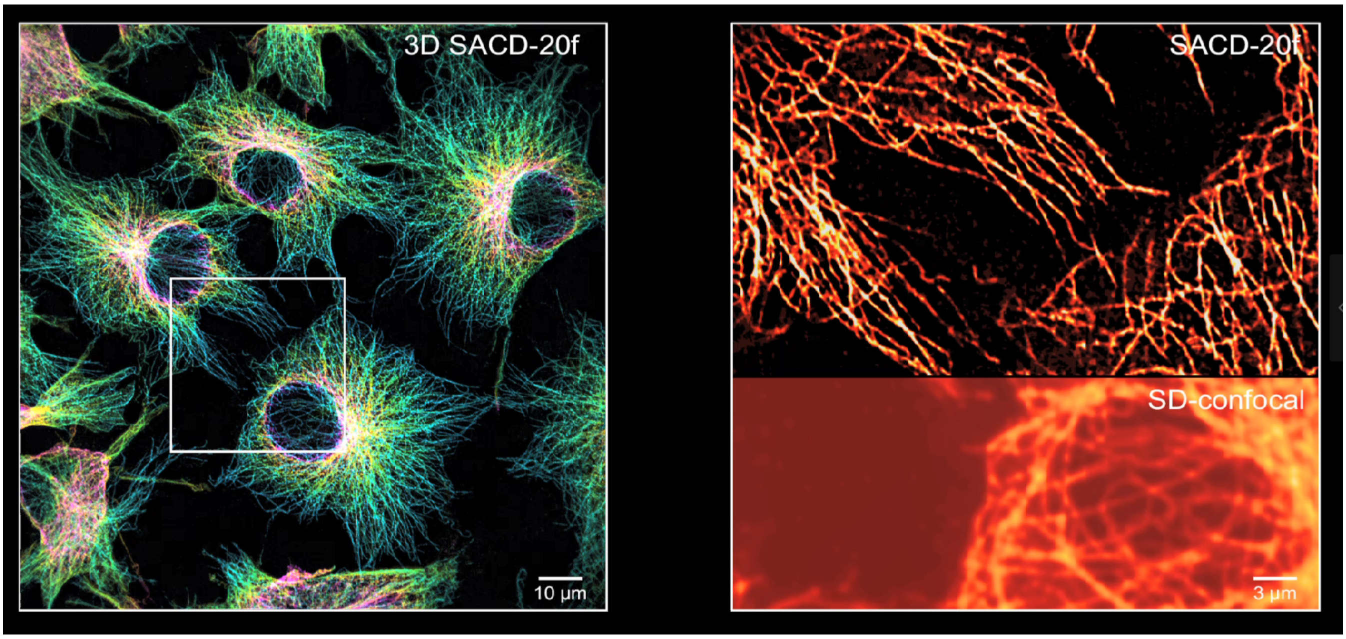


**Supplementary Video 1 | Large FOV SR imaging of 3D microtubule network.** Microtubule filaments in COS-7 cells labeled with $QD_{525}$ captured by SD-confocal and SACD-20f. (*c.f.*, **Fig. 2e**). Part I shows the color-coded projection of the microtubule filaments in the *z* direction as the depth increases gradually and the *y*-*z* orthoslices along the *x*-axis. Part II shows the magnified views of different regions from the whole FOV.

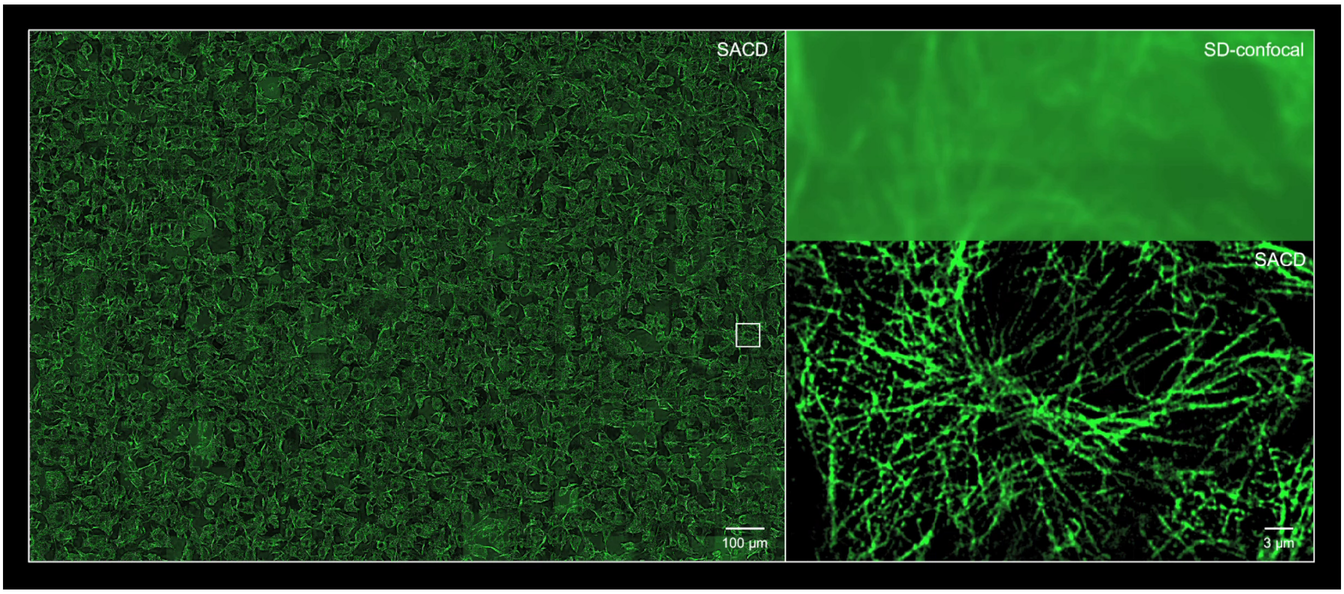


**Supplementary Video 2 | SACD enables high-throughput SR imaging.** Microtubule filaments in COS-7 cells labeled with $QD_{525}$ captured by SD-confocal and SACD-20f. (*c.f.*, **Fig. 3a**). Part I shows the microtubule filaments in a 2.0 mm × 1.4 mm area with 32 × 22 partly overlapping FOVs (66.6 μm × 66.6 μm each) in 20 minutes. Part II shows the magnified views of different regions from the whole FOV.

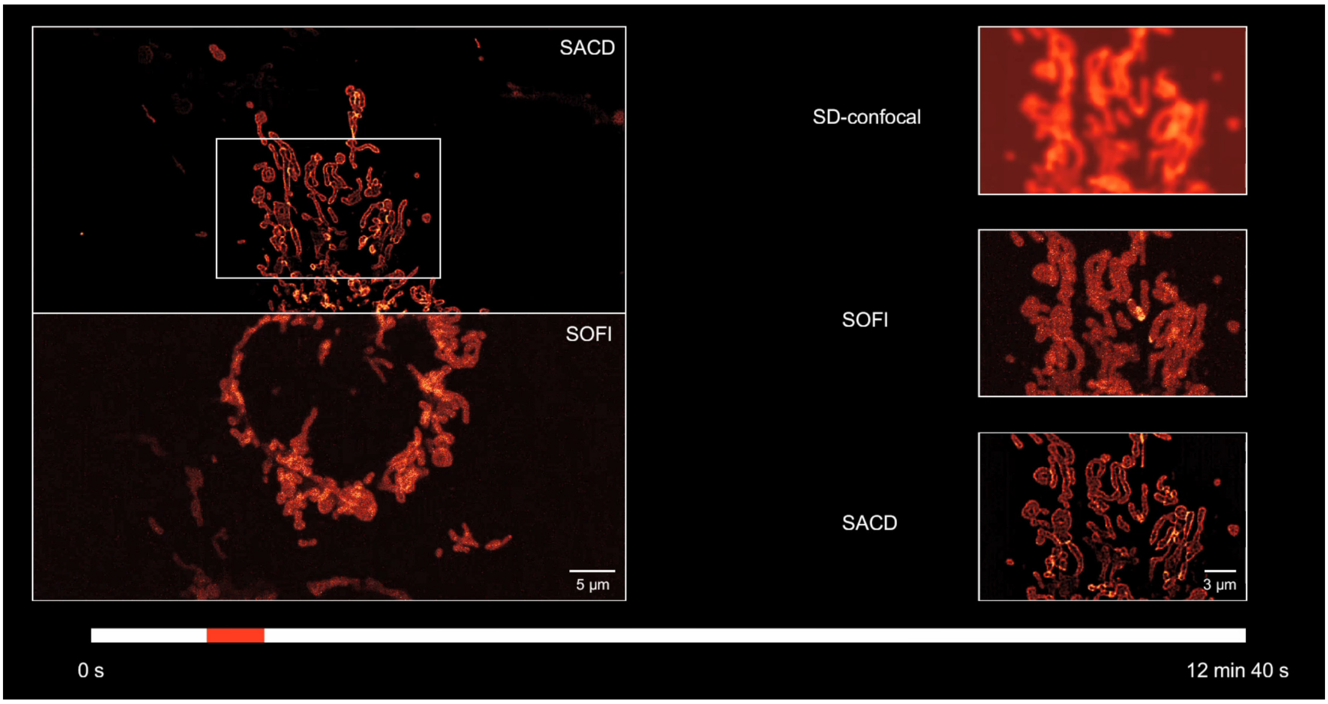


**Supplementary Video 3 | Comparisons of 20-frame SACD and 20-frame SOFI under live-cell conditions.** Part I demonstrates a representative live COS-7 cell labeled with Skylan-S-TOM20 imaged at 37°C by SD-confocal, SOFI-20f, and SACD-20f (*c.f.*, **Extended Data Fig. 9**). Part II compares the hollow outer mitochondrial membranes by SD-confocal, SOFI-20f, and SACD-20f for 20 time points at a 40 s interval.

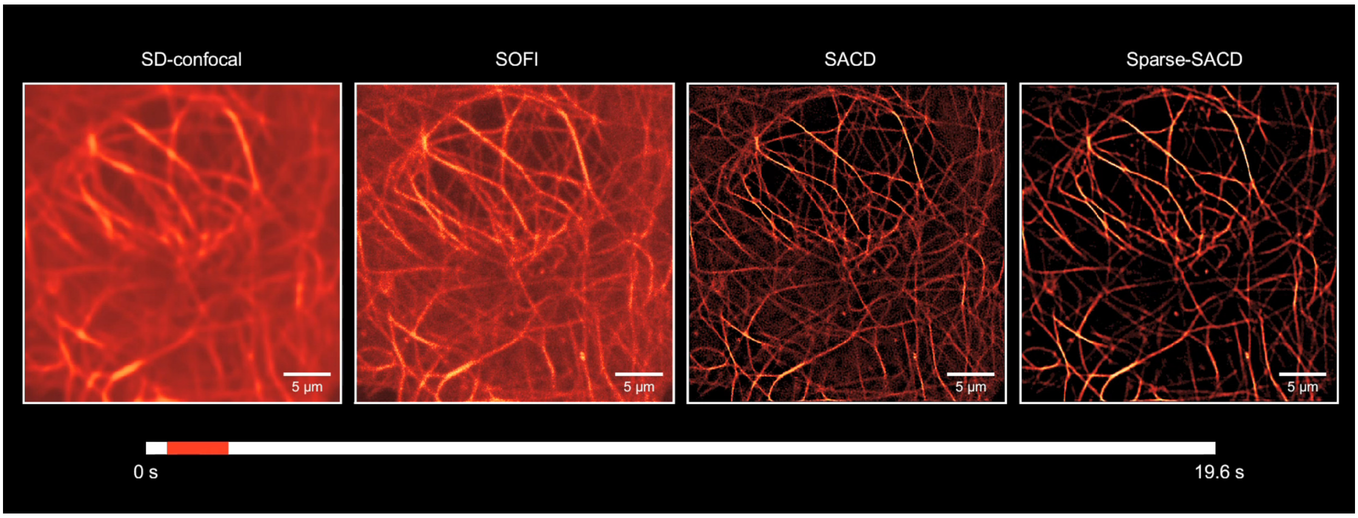


**Supplementary Video 4 | Continuous live-cell SR imaging of microtubules.** Part I demonstrates a representative live COS-7 cell labeled with MAP4-Skylan-S imaged at 37°C by SD-confocal, SOFI-20f, SACD-20f, and Sparse-SACD-20f (*c.f.*, **Fig. 4b**). Part II compares the microtubule filaments by SD-confocal, SOFI-20f, and SACD-20f for 50 time points at a 0.4 s acquisition time.

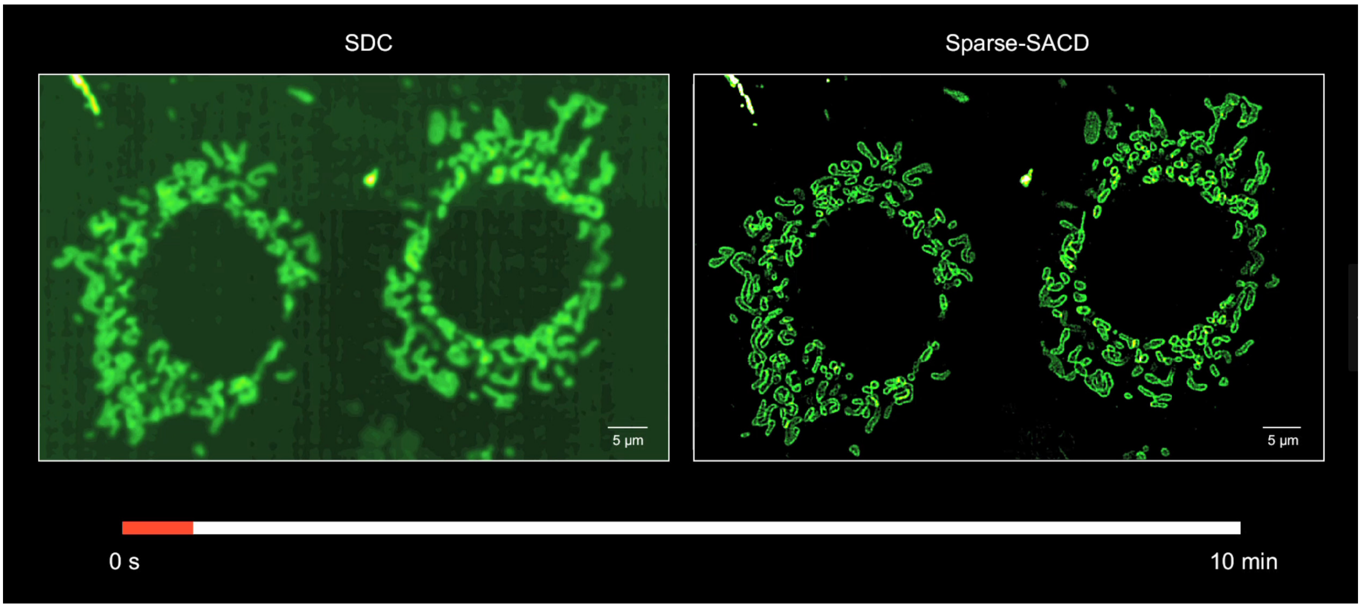


**Supplementary Video 5 | Long-term mitochondrial fission and fusion dissected by Sparse-SACD.** Part I demonstrates a representative live COS-7 cell labeled with Skylan-S-TOM20 imaged at 37°C by SD-confocal and Sparse-SACD-20f (*c.f.*, **Fig. 4d**). Part II shows the fission and fusion events detected by Sparse-SACD-20f for 16 time points at a 40 s interval.

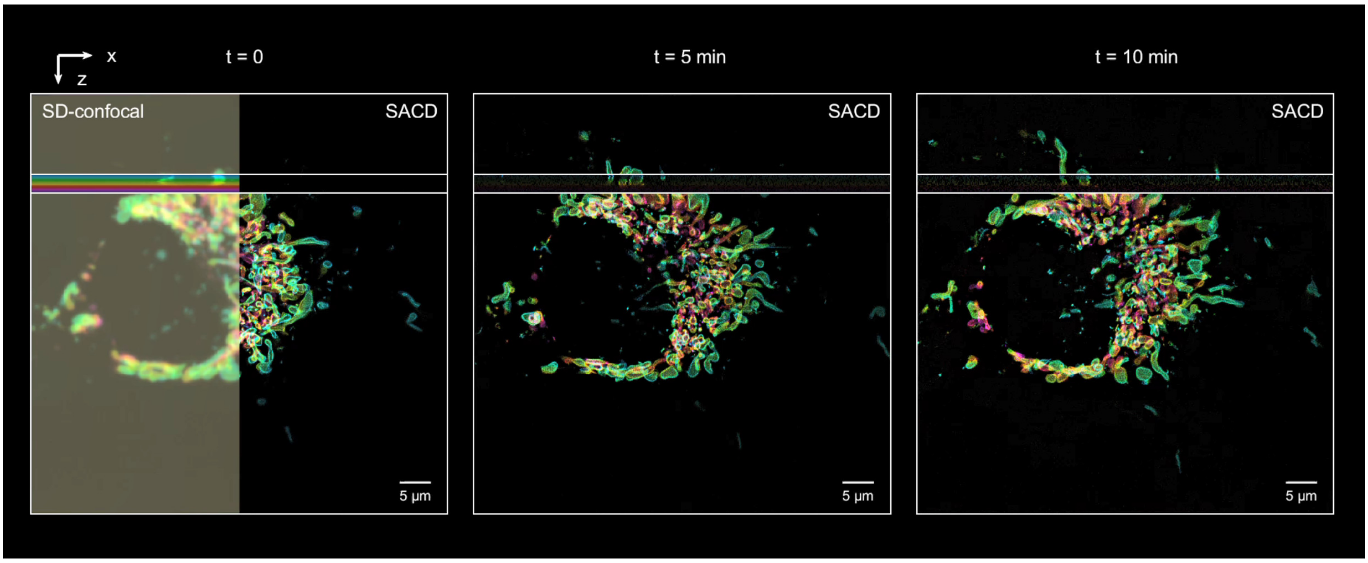


**Supplementary Video 6 | 4D SR imaging of mitochondrial network within a whole live cell across ten minutes.** Part I shows the comparison of SD-confocal and Sparse-SACD using a live COS-7 cell labeled with Skylan-S-TOM20 at 37°C (*c.f.*, **Fig. 4f**). Part II shows the color-coded projection of the hollow outer mitochondrial membranes in the *z* direction as the depth increases gradually and the *y*-*z* orthoslices along the *y*-axis.

#### References.
